# Eye metrics are a marker of visual conscious awareness and neural processing in cerebral blindness

**DOI:** 10.1101/2025.01.06.631506

**Authors:** Sharif I. Kronemer, Victoria E. Gobo, Shruti Japee, Eli Merriam, Benjamin Osborne, Peter A. Bandettini, Tina Liu

## Abstract

Damage to the primary visual pathway can cause vision loss. Some cerebrally blind people retain degraded vision or sensations and can perform visually guided behaviors. These cases motivate investigation and debate on blind field conscious awareness and linked residual neural processing. A key challenge in this research is that subjective measures of blind field visual conscious awareness can be misleading. Alternatively, eye metrics, including pupil size and eye movements are promising objective markers of conscious awareness and brain activity. In this study, we examined stimulus-evoked changes in pupil size, blinking, and microsaccades in the sighted and blind field of cerebrally blind participants. Using standard analysis and innovative machine learning methods, our findings support that eye metrics can infer blind field conscious awareness, even when behavioral performance on a visual perception task indicated otherwise. Furthermore, these eye metrics were linked to blind field visual stimulus-evoked occipital cortical field potentials. These findings support recording eye metrics in cerebral blindness and highlight potential clinical applications, including tracking the recovery of conscious vision and visual neural processing.

## Introduction

A person with cerebral blindness experiences partial or complete loss of conscious vision following a lesion to the visual pathway posterior to the lateral geniculate nuclei (e.g., optic radiations and primary visual cortex) ^1,2^. A longstanding question in cerebral blindness is whether visual neural processing persists, particularly in secondary visual pathways (e.g., the tectopulvinar pathway ^3,4^) and does any residual activity give rise to visual conscious awareness or behaviors typically associated with healthy conscious vision. Resolving these queries has significant implications for uncovering the function of the primary and secondary visual pathways in vision and behavior.

Motivating this curiosity are cerebrally blind people who can respond to visual stimulation and perform visually guided behaviors in their blind field, such as noticing visual movement, changes in environmental luminance, looking towards visual stimuli, or navigating an obstacle course ^5–7^. Similar findings are also reported in non-human primates with ablated primary visual cortex ^8–10^. For many cerebrally blind people, accurate blind field performance on visual tasks corresponds with visual conscious awareness, including degraded abnormal vision and non-visual “feelings” and “sensations” ^11–13^. However, a subgroup of cerebrally blind people is thought to have *blindsight*: the preservation of visually guided behaviors *without* visual conscious awareness ^14–17^. When instructed to perform a visual task, a person with blindsight will respond in the vein of “How can I look at something I haven’t seen?” ^6^. When told of their above chance performance, people with blindsight are often surprised because their actions felt random or by complete guessing ^4,6^.

Controversy persists whether people with blindsight genuinely experience total unconscious vision without any blind field visual conscious awareness ^18^. Partly fueling this debate are reports that residual conscious vision and degraded abnormal vision in cerebral blindness can be neglected by standard visual tasks and questionnaires (e.g., did you *see* or *not see* the image?) ^13,19^. Therefore, an objective measure of conscious perception may assist to probe blind field visual conscious awareness.

A promising marker of visual conscious awareness and linked residual neural processing in cerebral blindness is eye metric dynamics. In healthy physiology, eye metrics, including pupil size, blinking, and eye movements are linked with brain activity across species ^20–25^. Correspondingly, eye metrics are valuable to infer the consequences of brain activity, including those related to conscious awareness. For example, a previous study used pupil size, blink, and microsaccade responses to predict conscious perception of near-perceptual threshold stimuli ^22^. Furthermore, pupil size is indicative of conscious content, including the perceived or imagined brightness of an image, independent of physical luminance ^26–31^. Similarly, eye movements predict the direction of perceived motion ^32^.

In cerebral blindness, there is limited research on the relationship among eye metrics, conscious awareness, and neural processing. Among the available studies, pupillary changes are predominately examined. For example, several reports found preserved blind field pupillary light responses to ambient light and visual stimuli (e.g., grating images) ^11,33,34^. Likewise, pupillary responses to visual stimuli predict conscious vision and blindsight in cerebral blindness ^12,35^. Pupil size change is also responsive to complex visual features in healthy and cerebrally blind people. For instance, blind field pupillary responses (and facial muscle activity) were present for affective images of human facial expressions and body gestures ^36^. Meanwhile, less is known about blinking and eye movements in cerebral blindness. Previous studies find the maintenance of the blind field blink reflex, optokinetic nystagmus, and stereopsis ^6,37^, but there are conflicting results ^34,37^.

In the current experiment, we examined visual stimulus-evoked pupil size, blinking, and microsaccades in cerebral blindness. The visual stimuli were a curated set of images with physical and illusory attributes to help disentangle eye metrics linked with conscious versus unconscious neural processing. Alongside patients, we also tested age and education-matched control participants as a healthy comparison group. Finally, we recorded cortical activity with magnetencephalography (MEG) to relate behavior and eye metrics with cortical processing of visual stimuli.

We hypothesized that the control and patient participants would share similar visual stimulus-evoked pupil size, blink, and microsaccade dynamics for stimuli presented in the sighted field. Meanwhile, blind field eye metric dynamics would be present only for the physical but not illusory features of the visual stimuli. Likewise, we hypothesized that eye metrics would infer visual conscious awareness in the blind field, including the experience of residual and degraded conscious vision and non-visual sensations. These anticipated outcomes are significant for evidencing eye metrics as objective markers of conscious awareness and residual brain activity in response to visual stimulation in cerebrally blind people.

## Methods

### Participants

Eight cerebrally blind patient participants (females = 2; mean age = 50.25 years; standard deviation [SD] age = 22.76 years; mean education = 17.13 years; SD education = 2.59 years; Table 1) and eight age and education-matched healthy control participants (females = 4; mean age = 46.50 years; SD age = 18.69 years; mean education = 16.13 years; SD education = 1.89 years; Table 2) were recruited. The patient participants’ visual impairment consisted of left homonymous hemianopia (N = 2), right homonymous hemianopia (N = 2), left homonymous inferior quadrantanopia (N = 3), and right homonymous superior quadrantanopia (N = 1; Table 1; Supplementary Figure 1). Four additional patient participants were recruited but they were not included in analyses due to low data sample size, poor behavioral performance, and not meeting the definition of cerebral blindness (one recruited patient participant experienced visual impairment due to a chiasmal tumor).

**Table 1.**
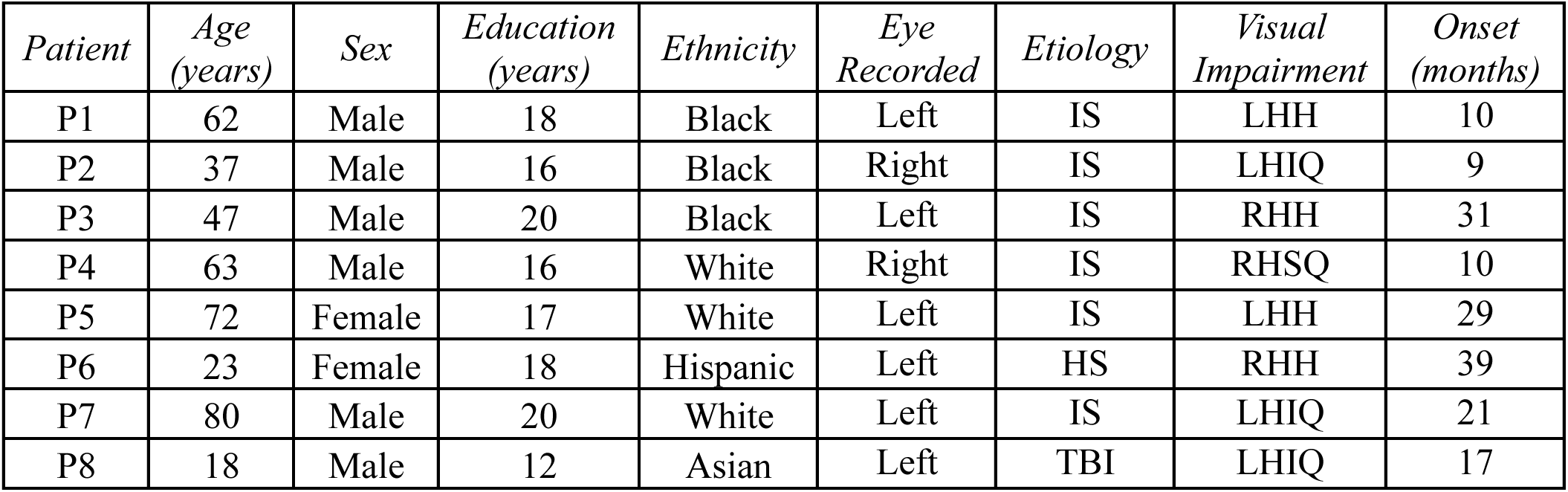
Patient participants’ demographic information (age, sex, education, and ethnicity), eye recorded, etiology, visual impairment, and duration since injury onset. *Etiology*: Ischemic stroke (IS); hemorrhagic stroke (HS); traumatic brain injury (TBI). *Visual impairment*: Left homonymous hemianopia (LHH); right homonymous hemianopia (RHH); left homonymous inferior quadrantanopia (LHIQ); right homonymous superior quadrantanopia (RHSQ).

| *Patient* | *Age*<br>*(years)* | *Sex* | *Education*<br>*(years)* | *Ethnicity* | *Eye*<br>*Recorded* | *Etiology* | *Visual*<br>*Impairment* | *Onset*<br>*(months)* |
| --- | --- | --- | --- | --- | --- | --- | --- | --- |
| P1 | 62 | Male | 18 | Black | Left | IS | LHH | 10 |
| P2 | 37 | Male | 16 | Black | Right | IS | LHIQ | 9 |
| P3 | 47 | Male | 20 | Black | Left | IS | RHH | 31 |
| P4 | 63 | Male | 16 | White | Right | IS | RHSQ | 10 |
| P5 | 72 | Female | 17 | White | Left | IS | LHH | 29 |
| P6 | 23 | Female | 18 | Hispanic | Left | HS | RHH | 39 |
| P7 | 80 | Male | 20 | White | Left | IS | LHIQ | 21 |
| P8 | 18 | Male | 12 | Asian | Left | TBI | LHIQ | 17 |

**Table 2.** Control participants’ demographic information (age, sex, education, and ethnicity), eye recorded, and their paired patient participants (see Table 1).

| *Control* | *Age*<br>*(years)* | *Sex* | *Education*<br>*(years)* | *Ethnicity* | *Eye Recorded* | *Paired*<br>*Patient* |
| --- | --- | --- | --- | --- | --- | --- |
| C1 | 63 | Female | 17 | White | Right | P1 |
| C2 | 29 | Male | 16 | White | Right | P2 |
| C3 | 50 | Male | 16 | Hispanic | Right | P3 |
| C4 | 62 | Female | 16 | White | Left | P4 |
| C5 | 59 | Male | 18 | White | Left | P5 |
| C6 | 23 | Female | 16 | Biracial | Left | P6 |
| C7 | 64 | Female | 18 | White | Left | P7 |
| C8 | 22 | Male | 12 | White | Left | P8 |

The patient participants were recruited from the National Institute of Neurological Disorders and Stroke in Bethesda, Maryland (MD), USA and MedStar Georgetown University Hospital in Washington, District of Columbia, USA. Control participants were recruited from the local Bethesda, MD, USA community. All participants were recruited, consented, and tested in accordance with protocols approved by the Institutional Review Board of the National Institute of Mental Health.

All patient participants completed two behavioral study sessions, except for participant P6 who completed only one session because they were lost to follow-up. P4 completed an additional magnetencephalography (MEG) study session (see *Magnetencephalography* section). All control participants completed one behavioral study session. Less total trials were required for control participants because the left and right visual field stimulus presentation locations (see *Visual Perception Task* section) were combined in analyses, as both locations were sighted in control participants.

*Inclusion criteria* included: (1) 18 years of age or older, (2) at least a high school education (12 years or more), (3) capacity to provide their own informed consent, and understand and cooperate with study procedures, (4) neurologically normal (control participants only), and (5) unilateral or bilateral focal lesions and at least three months post-lesion (patient participants only). The average injury duration relative to the date of participation in the current study was approximately 21 months (Table 1).

*Exclusion criteria* included: (1) any neurological or psychiatric disorder unrelated to the focal lesion (e.g., epilepsy and schizophrenia; patient participants only), (2) previous head injury (control participants only), (3) present or past (within six months) drug or alcohol abuse or addiction, and (4) radiation treatment to the brain during a three-month period prior to the experiment (patient participants only).

The cause of cerebral blindness in the patient participants included ischemic stroke, hemorrhagic stroke, and traumatic brain injury (Table 1). Strokes were related to hypertension, diabetes, atrial fibrillation, vascular surgery, a skiing accident, and a fall. For example, P3 suffered an ischemic stroke in the left cortical hemisphere. A structural whole brain magnetic resonance imaging (MRI) scan acquired at the time of the study revealed a lesion that includes the left optic radiation, fusiform gyrus, and primary visual cortex (see *Structural Magnetic Resonance Imaging* section; Figure 1A). A Humphrey visual field (HVF) test obtained approximately 18 months before the first study session found that P3 was impaired with right homonymous hemianopia (Figure 1B; Supplementary Figure 1). See Table 1 and Supplementary Figure 1 for clinical details and HVF test results for all patient participants.

**Figure 1.**
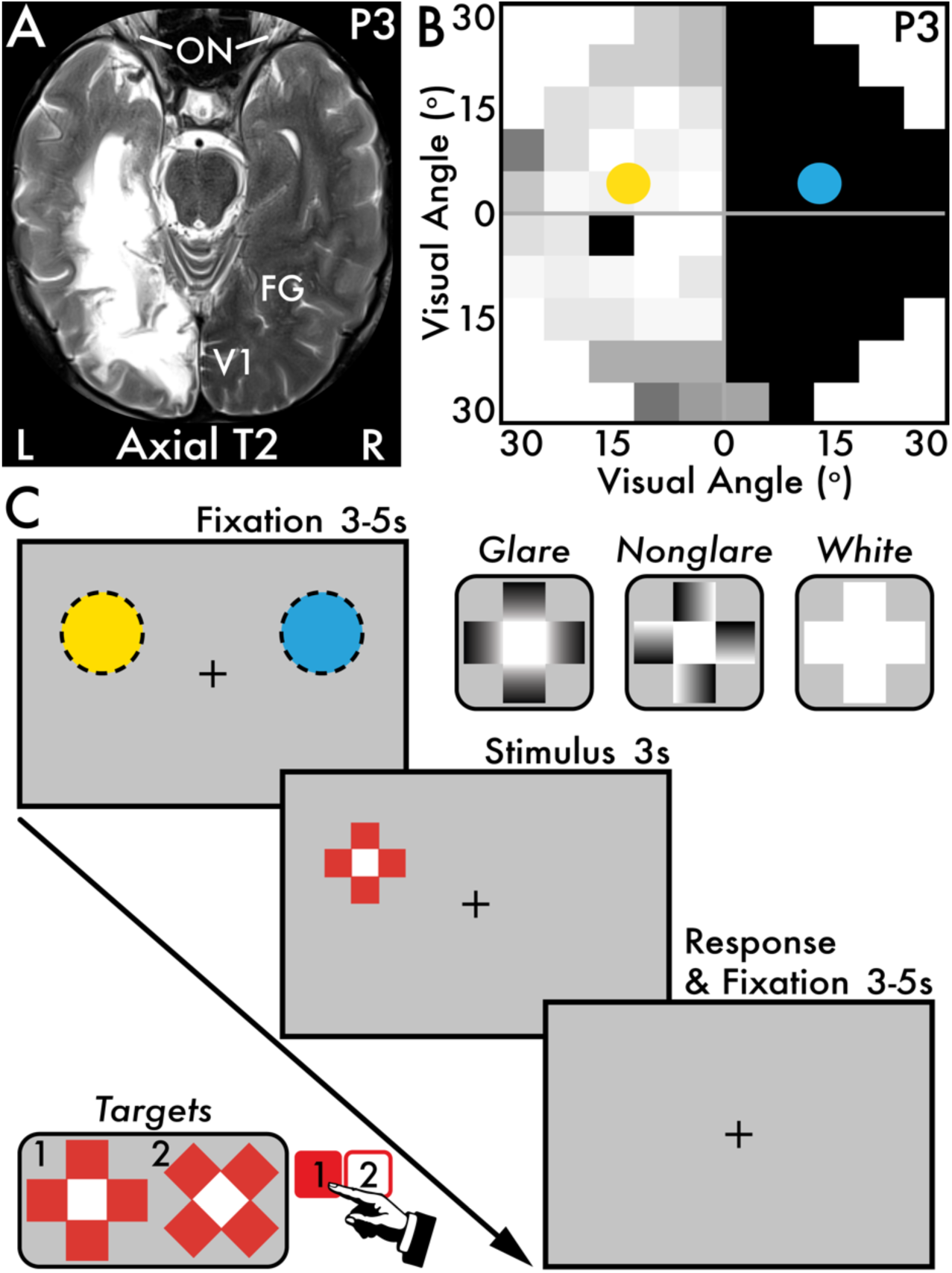
Patient participant P3 stroke location and Humphrey visual field test results, and the visual perception task. **(A)** Patient participant P3 axial T2-weighted whole brain MRI scan revealing a left (L) cortical hemisphere stroke affecting the left optic radiation, fusiform gyrus (FG), and primary visual cortex (V1). The right (R) cortical hemisphere is intact. Major anatomical landmarks are labeled: optic nerve (ON), FG, and V1. **(B)** P3’s Humphrey visual field (HVF) test results from the left eye in degrees (°) of visual angle. Sighted areas are indicated by white and light gray squares, while blind areas are indicated by dark gray and black squares. The single black square in the left visual field is the natural blind spot. The colored circles (sighed field: yellow; blind field: blue) indicate the on-screen stimulus presentation locations of the visual perception task. **(C)** Each visual perception task trial involved three main phases: (1) a pre-stimulus fixation period (3-5 seconds [s]), (2) a stimulus presentation period (3 s), and (3) a post-stimulus response and fixation period (3-5 s). Throughout all phases in a trial, participants were instructed to maintain fixation on the central plus sign image. There were two categories of stimuli: (1) target and (2) nontarget. The target stimuli were red with a white center and appeared in equal proportion between two orientations: (1) plus sign and (2) x-oriented. Participants were instructed to make an immediate keypress upon perceiving the target stimulus and indicate which orientation the target appeared (e.g., 1-key = plus sign-oriented; 2-key = x-oriented). The key mapping to target stimulus orientation was counterbalanced across participants. There were three nontarget stimuli that appeared in equal proportion: (1) glare, (2) nonglare, and (3) white. Participants were instructed that no keypress was required when the nontarget stimuli appeared. The yellow (sighted) and blue (blind) dotted circles approximate the stimulus presentation locations for P3, corresponding with the circles overlaid on the HVF test results in B. The stimulus presentation locations were adjusted for each patient participant according to their visual field impairment (Table 1; Supplementary Figure 1). Control participants were shown stimuli in the exact same on-screen locations as their paired patient participants (Table 2). Stimuli appeared in equal proportion between the two stimulus presentation locations.

### Visual Perception Task

The visual perception task consisted of trials with three primary phases (Figure 1C): (1) a pre-stimulus fixation period (jittered 3-5 seconds), (2) a stimulus presentation period (3 seconds), and (3) a post-stimulus response and fixation period (jittered 3-5 seconds). Across all task phases, a central fixation cross (a black plus sign; behavioral session: visual angle = 0.99 x 0.99 degrees; MEG session: visual angle = 0.38 x 0.38 degrees) continuously appeared on a blank, gray screen. For patient participants P7 and P8 and their paired control participants C7 and C8, the fixation cross was positioned to the top-center of the behavioral display monitor (see *Equipment and Testing Facility* section) because the blind field for these patient participants began deeper in their peripheral vision, inaccessible to an on-screen stimulus presentation location with centrally fixation. Control and patient participants completed approximately 10 seven-minute task blocks comprising 40 trials each. After each task block, participants were prompted to share by verbal report whether they saw or felt the presence of any task stimuli in their sighted and blind field (see Table 3 and 4).

**Table 3.** Summary of *blind aware* patient participants’ blind field task behavior, eye metrics, and verbal perceptual report results (see *Correspondence among blind field task behavior, verbal perceptual report, and eye metric dynamics* Methods section). *Patient participant P4 (Table 1) task behavior was absent in the original visual perception task but present in the behavioral-adapted task (Figure 5B versus C).

| Blind Aware |  |  |  |  |
| --- | --- | --- | --- | --- |
| *Patients* | *Task Behavior* | *Eye Metrics* | *Verbal Report* | *Verbal Report Summary* |
| P2 | Present | Present | Present | <u>Summary:</u> Reported seeing or sensing most task stimuli.<br><u>Representative quotes:</u> “I can see it [images] but not processing”; “I can feel something is there but cannot tell you where it is or what it is”; “the left [blind field] images were incomplete like a bad signal”; “takes time to perceive them and figure out the shape”; “barely could see images on the left [blind field]”; “images kind of obscured – not in focus – on the left [blind field] but could see something was there”. |
| P4 | Present* | Present | Present | <u>Summary:</u> Reported seeing or sensing most task stimuli.<br><u>Representative quotes:</u> “I could sense there was movement in the upper right [blind field] but could not see anything”; “feeling of movement [or] clicking done in the upper right quadrant [blind field] ... but no vision”; “[blind field looked like] static noise; old time TV”; “feels like an area with static noise would be added when there was not any before”; “[had sensation of images akin to movement but] cannot see what it is”; “[sensation of images were] like a vibration”; “strong feeling [of an image]”. |
| P7 | Present | Present | Present | <u>Summary:</u> Reported seeing most task stimuli.<br><u>Representative quotes:</u> “perhaps [images] more faded and lacked color”; “[the images were] a little more washed out”; “[images] just did not look as sharp on the left [blind field]”; “[images have] more contrast and brighter on the right [sighted field]”; “confident about color but not shape [of images in the blind field]”; “more aware of images on the right [sighted field]”. |
| P8 | Absent | Present | Present | <u>Summary:</u> Reported seeing most task stimuli.<br><u>Representative quotes:</u> “felt there was a fog over the left side [blind field]”; “could not tell the shape [of images]”; “not sure if I mistook a gray for red [image]”; “barely only able to catch the left [blind field] images”; “left [blind field] images were smudged”; “could catch a few on the left [blind field] but cannot really tell”; “surprised that when I looked on the left [blind field] that there was a red image – thought it was black”; “on the left [blind field] what might be red is grayish”. |

Following the initial fixation period, a visual stimulus appeared (behavioral session: visual angle = 6.51 x 6.51 degrees; MEG session: visual angle = 5.19 x 5.19 degrees). Participants were instructed to always maintain their gaze on the fixation cross and never to directly look at the stimuli that appeared in the periphery of the screen. The stimuli could appear in one of two mirroring on-screen locations equal distant from fixation. For the patient participants, one stimulus location was positioned in their sighted field and the second was positioned in their blind field. During each task trial, a single stimulus would appear in either on-screen location at random but in equal proportion within task blocks.

The sighted and blind field locations were determined in a pre-task stimulus location calibration phase that was guided by the patient participants’ HVF test results that were made available to the experimenters prior to each study session, except for P6 who did not complete a HVF test in time for their study session (Supplementary Figure 1). During location calibration, a central fixation and two nonglare stimuli (see stimulus details below) appeared in mirroring on-screen locations. Next, the experimenter manually adjusted the stimuli locations while the patient participants centrally fixated and indicated by verbal report their conscious awareness for the on-screen stimuli. The stimulus presentation locations were set when the patient participant reported that they perceived only the stimulus positioned in their sighted field, while no longer able to see or sense the stimulus positioned in their blind field. For P2 and P7, there was no blind field, on-screen location where these participants reported *no* conscious awareness of the stimulus during location calibration. The colored circles overlaid on the HVF test in Figure 1B and Supplementary Figure 1 approximates the stimulus size and the sighted (yellow; P4 MEG session: orange) and blind field (blue; P4 MEG session: green) stimulus presentation locations for each patient participant. For control participants, the location calibration phase was not completed. Instead, the same stimulus presentation locations were used as their paired patient participants (Table 2).

The visual perception task presented five visual stimuli: (1) glare, (2) nonglare, (3) isoluminant, (4) white, and (5) red (Figure 1C; see Supplementary Figure 2A for the isoluminant stimulus). Each stimulus type was shown in equal proportion within task blocks and between stimulus presentation locations but in random order. There were two main stimulus categories: (1) target and (2) nontarget.

The target was the red stimulus. In an early version of the task (administered to participants P1 session 1, C1, and C2), there was only one type of target stimulus: a plus sign-oriented image with a central white square and four surrounding red squares (Figure 1C). The remaining participants and P1 session 2 completed an updated task version with *two* target stimulus types that appeared in equal proportion: (1) plus sign and (2) x-oriented (Figure 1C). Both target stimulus types were identical except the x-oriented target stimulus was rotated 45 degrees relative to the plus sign-oriented target stimulus.

The two-target task version was introduced to test the visual acuity of patient participants who reported conscious awareness for task stimuli in their blind field. In the one-target task version, participants were prompted to select a single key immediately upon perceiving the target stimulus, no matter its on-screen presentation location. In the two-target task version, participants were prompted to select one of two keys upon perceiving the target stimulus, no matter its on-screen presentation location. Each key corresponded with either the plus sign or x-oriented target stimulus. The key mapping with the target orientation type was counterbalanced across participants.

The nontarget stimuli consisted of the glare, nonglare, isoluminant, and white stimulus. The glare stimulus was a plus sign-oriented image with a central white square and four surrounding squares with a black-to-white gradient facing inward (i.e., the white portion of the gradient contacted the central white square). The glare stimulus and similar versions of this image are reported to induce the illusory perception of glare or brightness ^38,39^. The nonglare stimulus was identical to the glare stimulus except the surrounding black-to-white gradient squares were oriented along the central white square and did not induce the perception of illusory brightness (Supplementary Figure 2B). The isoluminant stimulus was identical to the nonglare stimulus except the central white square was replaced with a gray square that exactly matched the gray background of the screen on which all stimuli appeared. Also, the black-to-white gradient squares in the glare, nonglare, and isoluminant stimulus had an average luminance equal to the gray screen background. Therefore, the average total luminance of the isoluminant stimulus was equal to the gray screen background. Eye metric results for the isoluminant stimulus are not shown. Finally, the white stimulus appeared as an all-white version of the glare, nonglare, and isoluminant stimulus. For all nontarget stimuli, participants were instructed to withhold any response when they appeared (i.e., nontarget stimuli were task irrelevant).

For patient participant P4, the visual perception task was modified into two additional task versions: (1) behavioral and (2) MEG-adapted. The behavioral-adapted visual perception task maintained the identical trial structure as the original visual perception task (Figure 1C). However, the behavioral-adapted task only presented the target stimulus (i.e., plus sign and x-oriented red stimuli). P4 was instructed to make an immediate keypress whenever he perceived the target stimulus in his sighted field or experienced what he described as a vision or feeling of “movement” and “vibration” in his blind field (see *Visual perception task* Visual behavior Results section; Table 3). Similar to the original perception task, P4 was instructed to make a keypress to indicate the target orientation and, if uncertain, to make his best guess for its orientation. The stimulus presentation locations were identical to those used for P4 in the original visual perception task (Supplementary Figure 1).

The MEG-adapted visual perception task maintained the trial structure as the original visual perception task (Figure 1C). However, this task version only presented the target stimuli and a subset of the nontarget stimuli: (1) glare and (2) nonglare. This adaptation was made to increase the number of trials for these stimulus types. Also, because P4 suffered from right homonymous superior quadrantanopia (upper right quadrant vision loss) the sighted stimulus position was adjusted from the behavioral task to appear in the sighted, bottom right quadrant of the visual field (Supplementary Figure 1). This adjustment was made so that the MEG field potentials for stimuli presented in the sighted and blind field, now both located in the right visual field, would correspond with contralateral field potential changes in the *left* visual cortex.

### Brightness Perception Task

Previous studies report that the glare stimulus is perceived with illusory brightness ^39^. To gauge if the participants in the current experiment also perceived illusory brightness from the glare stimulus, participants were asked to complete a brightness perception task (Supplementary Figure 2A). Each task trial began with a blank gray screen (2 seconds). Next, participants were shown dual combinations of the glare, nonglare, and isoluminant stimulus (see *Visual Perception Task* section for stimulus details) and instructed to report with keypresses which stimulus appeared brighter near its center or if both images appeared with equal brightness. Participants could look at the images directly and their responses were self-paced. Participants completed a single 30-trial task block with 10 trials each of the following stimulus comparisons: (1) glare versus nonglare, (2) glare versus isoluminant, and (3) nonglare versus isoluminant. The on-screen stimulus presentation locations were identical for all participants and the patient participants reported being able to clearly see the stimuli prior to making a response.

### Pupillometry and Eye Tracking

Head-fixed (SR Research Head Support; SR Research, Inc.) monocular pupillometry and eye tracking were acquired with the EyeLink 1000 Plus (sampling rate = 1000 Hz; SR Research, Inc.). Whichever eye was best positioned with the eye tracker camera was selected for recording (Table 1 and 2). The EyeLink 1000 Plus software and monitoring of eye tracking during the study session was performed on a Dell OptiPlex XE2 desktop computer and monitor (Dell, Inc.). The behavioral computer (see *Testing Equipment and Facility* section) and EyeLink desktop communicated via an Ethernet connection. Participants were positioned approximately 56 and 106 centimeters from the EyeLink camera in the behavioral and MEG study sessions, respectively.

### Magnetencephalography

MEG data were recorded using a CTF 275 MEG system (sampling rate = 1200 Hz; CTF Systems, Inc., Canada) composed of a whole-head array of 275 radial 1st order gradiometer sensors housed in a magnetically shielded room (Vacuumschmelze GmbH & Co. KG, Germany). Data were not recorded from three malfunctioning sensors, including one right occipital sensor (O13; Supplementary Figure 7B). Synthetic 3rd gradient balancing was used to remove background noise. The MEG acquisition software was run on a Dell Precision T7500 desktop computer (Dell, Inc.). While recording the MEG data, P4 completed the MEG-adapted visual perception task (see *Visual Perception Task* section). Behavioral task events (e.g., stimulus onset) were marked in the MEG recording via parallel port. Simultaneously, head-fixed pupillometry and eye tracking were recorded with the EyeLink 1000 Plus system (SR Research, Inc.; see *Pupillometry and Eye Tracking* section).

### Structural Magnetic Resonance Imaging

A whole brain structural MRI was acquired on a 3T MAGNETOM Skyra MRI (Siemens, Inc.; axial T2-weighted image: repetition time = 3.5 seconds; echo time = 0.086 seconds; flip angle = 120 degrees).

### Testing Equipment and Facility

All experimental sessions were completed in a windowless, temperature-controlled room at the National Institutes of Health, Bethesda, MD, USA. Each study session lasted approximately 2.5 hours, including a health exam, instructions, and task breaks. During the behavioral session, the experimenters were positioned behind the participant to monitor behavior and deliver task instructions. During the MEG session, the experimenters were positioned outside the MEG shielded room and monitored and communicated with the participant via a closed-circuit television (COLOR CCD Camera VCC-3912; Sanyo Electric Co. Ltd.) and intercom console system (VSM MedTech Ltd.).

The behavioral tasks (see *Visual Perception Task* and *Brightness Perception Task* sections) were coded in Python and run with PsychoPy (behavioral session: v2022.2.4; MEG session: v2021.1.2; Open Science Tools Ltd.). During the behavioral study sessions, the tasks were run on a behavioral laptop (MacBook Pro 2019; 13-inch; 2560 x 1600 pixels; Mac OS Catalina v10.15.7; Apple, Inc.) and participants viewed the behavioral tasks on a VPixx monitor (1920 x 1200 pixels; VPixx Technologies, Inc.) that mirrored the laptop display via a DVI connection. During the MEG study session, the behavioral task was run on a Dell Precision T3500 desktop computer (Dell, Inc.). Patient participant P4 viewed the behavioral task on a projector screen that mirrored the desktop display via a PROPixx LED projector (VPixx Technologies, Inc.) positioned outside the MEG shielded room. The projected light first passed through a waveguide, then an adjustable two-mirror system that reflected the display image onto the projection screen. Participants were positioned approximately 58 and 75 centimeters from the behavioral monitor and MEG projector screen, respectively.

For all study sessions, participants were instructed to make their responses during the behavioral task with their right hand. During the behavioral study session, participants made keypresses using a keyboard positioned on a table in front of the participant. During the MEG study session, P4 made button presses using a button box (4 Button Inline Fiber Optic Response Pad; Current Designs, Inc.). Button presses were received via an electronic interface (932 Interface & Power Supply; Current Designs, Inc.) and marked in dedicated channels of the MEG recordings.

### Statistical Analysis

All analyses were completed in MATLAB (R2023b; Mathworks, Inc.). Data and results were visualized using MRIcron (https://www.nitrc.org/projects/mricron), MATLAB (R2023b; Mathworks, Inc.), Prism (version 10.4.0; GraphPad Software, LLC.), and Illustrator (Adobe, Inc.).

### Demographics

Control versus patient participant age and education were statistically compared with an independent-samples *t*-test.

### Visual perception task

Perception rate and orientation discrimination accuracy rate were calculated for the target stimulus (see *Visual Perception Task* section; Figure 2; Figure 5B, C, and E). Note that perception rate and accuracy rate could not be calculated for the nontarget stimuli because participants did not respond to these stimuli (i.e., nontarget stimuli were task irrelevant). *Perception rate* was calculated as the number of *perceived* target stimuli divided by the total number of *presented* (perceived + not perceived) target stimuli. A target stimulus was considered perceived if the participant responded with a key or button press within 5000 milliseconds from the onset of the target stimulus, regardless of orientation accuracy. *Accuracy rate* was calculated as the number of *correct perceived* target stimulus orientation type (see *Visual Perception Task* section) divided by the total number of *all perceived* (correct + incorrect) target stimuli. Chance level target orientation accuracy rate was 0.5.

**Figure 2.**
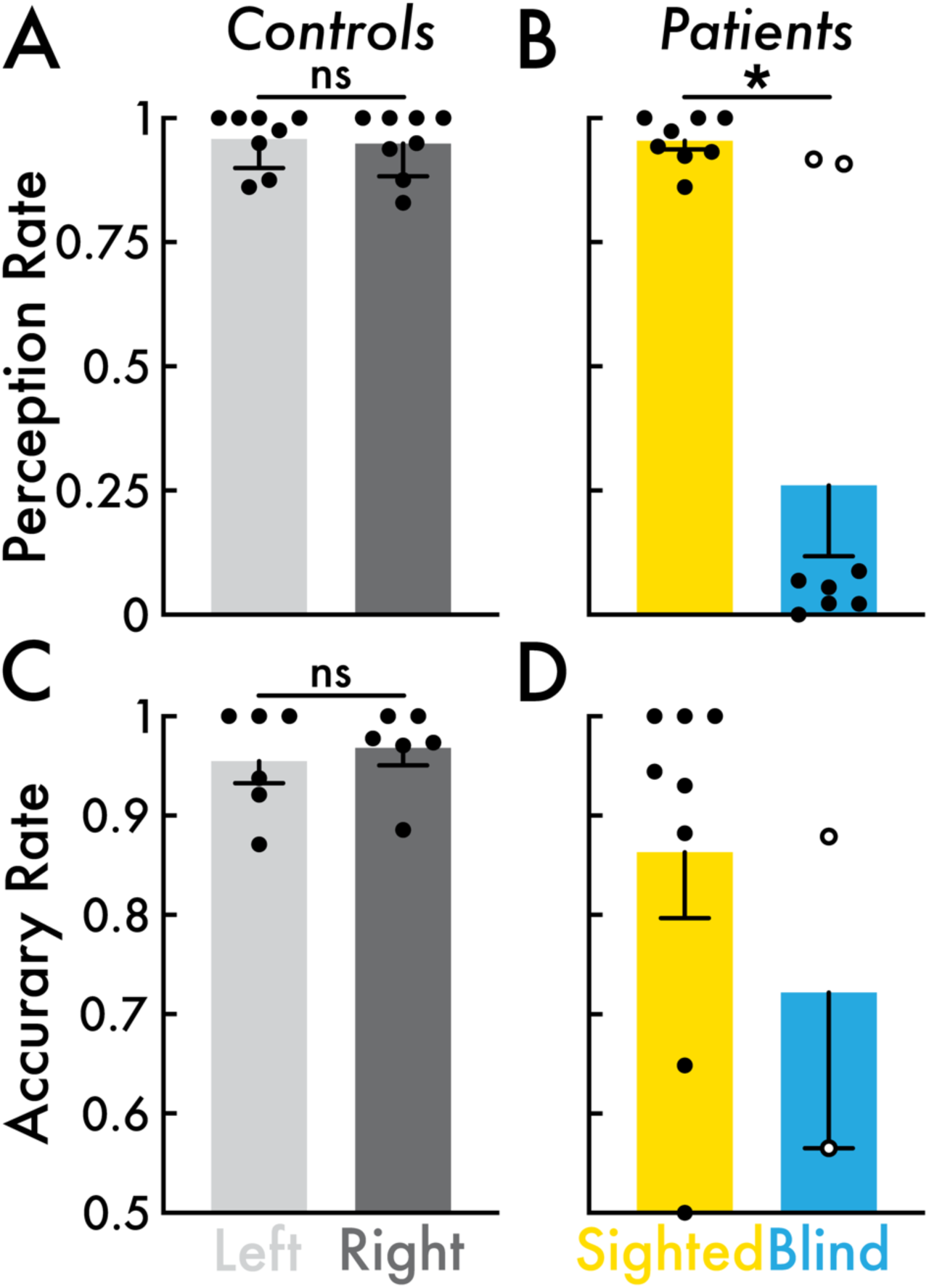
Visual perception task behavioral results. **(A)** Control participants (N = 8) target stimulus perception rate for the left (light gray) and right (dark gray) visual field. Left versus right visual field perception rates were not significantly different (ns). **(B)** Patient participants (N = 8) target stimulus perception rate for the sighted (yellow) and blind field (blue). Open circles highlight patient participants P2 and P7 who reported high (> 0.9) target stimulus perception rate in their blind field. Sighted versus blind field perception rates were significantly different (*; *p* < 0.05). **(C)** Control participants (N = 6) target stimulus orientation discrimination accuracy rate for the left and right visual field. Accuracy rate could not be calculated for control participants C1 and C2 (Table 2) who completed an early version of the visual perception task (see *Visual Perception Task* Methods section) that did not include a target orientation discrimination component. Left versus right visual field accuracy rates were not significantly different. **(D)** Patient participants target stimulus orientation discrimination accuracy rate for the sighted (N = 8) and blind (N = 2) field. Blind field accuracy rate was only calculated for patient participants P2 and P7 with high blind field perception rate (highlighted with open circles in B). Sighted versus blind field accuracy rate was not statistically evaluated. In all subplots, the open and closed circles represent individual participants, and the bars and error bars indicate the group mean and standard error of the mean, respectively.

For control participants, perception rate and accuracy rate were calculated separately within the *left* and *right* visual field stimulus presentation locations (Figure 2A and C). For patient participants, perception rate and accuracy rate were calculated separately within the *sighted* and *blind* field stimulus presentation locations (Figure 2B and D). Accuracy rate could not be calculated for C1 and C2 who completed the one-target visual perception task version (see *Visual Perception Task* section). Blind field accuracy rate was calculated for only patient participants P2 and P7 who reported high (> 0.9) blind field perception rate (open circles in Figure 2B; see *Behavior* Results section). The remaining patient participants reported low (< 0.1) blind field perception rate (closed circles in Figure 2B; see *Behavior* Results section), thus too few perceived target stimulus trials to reliably calculate accuracy rate. Left versus right (control participants) and sighted versus blind (patient participants) perception rate and accuracy rate were statistically compared with a paired *t*-test.

Additional behavioral analyses were evaluated for P4’s performance on the behavioral-adapted visual perception task. First, the false positive rate was calculated as the number of key presses *more than* 5000 milliseconds from the onset of a target stimulus divided by the total number of key presses. Also, P4’s keypress reaction times relative to stimulus onset were statistically compared between the sighted and blind field with a paired *t*-test. Finally, to assess whether P4’s target orientation accuracy rate was statistically greater than chance, a one-sided binomial test was conducted.

### Brightness perception task

Perceived brightness of the glare, nonglare, and isoluminant stimulus was calculated using a custom scoring system. For each trial that the participant reported a stimulus as brighter than the juxtaposed image, the stimulus perceived as brighter received 1 point; 0.5 points if the stimulus was reported as equally bright; and 0 points if the stimulus was reported as less bright. Participants completed 10 trials each of the following stimulus comparisons: (1) glare versus nonglare, (2) glare versus isoluminant, and (3) nonglare versus isoluminant. Thereby, a stimulus that was always reported as brighter would receive a maximum score of 20, while a stimulus that was never reported as brighter would receive a minimum score of 0. The brightness perception reports were tested with a one-way ANOVA and post-hoc paired *t*-test (glare versus nonglare and nonglare versus isoluminant; Holm-Bonferroni corrected for multiple comparisons; Supplementary Figure 2B and C).

### Eye metrics

#### Pupil size epoch extraction

The pupil data were preprocessed, including the removal of blinks (*stublinks.m*; available at http://www.pitt.edu/~gsiegle; ^40^) and smoothing. The preprocessed pupil data were cut into 18001-millisecond epochs centered at stimulus onset. Also, interstimulus interval or blank event epochs were cut for each stimulus type, centered between 4000 and 7000 milliseconds after stimulus onset (i.e., a minimum of 1000 milliseconds after the offset of the preceding stimulus and before the onset of the subsequent stimulus). The blank event time was calculated by selecting a random time within the interstimulus interval following each stimulus and preceding the subsequent stimulus or task block end time (for the final trial blank event in each task block). Therefore, each stimulus epoch had a corresponding blank event epoch. Finally, all stimulus and blank event epochs were baselined to the average pupil size within the 1000 milliseconds immediately preceding the stimulus onset or blank event. Pupil size epochs were removed from analysis if between 1000 milliseconds pre-stimulus onset or blank event and 6000 milliseconds post-stimulus onset or blank event there was an extreme pupil size value (> 1750 pixels; this value was selected by visual inspection of the pupil size epoch timecourses) or if more than 50 percent of the samples within this epoch interval did not have a pupil size value (e.g., due to prolonged eye closure or loss of eye tracking). Across all subjects and events, these exclusion criteria resulted in the removal of less than 15 percent of pupil size epochs. Finally, epochs were averaged within participant across event type (e.g., the nontarget condition results were an average of the glare, nonglare, and white stimulus epochs) and stimulus location.

#### Blink epoch extraction

Blinks were determined by the pupil size preprocessing method that identified blink intervals (see *Pupil size epoch extraction* section). The resulting blink data set was a binary timecourse (0 = blink absent; 1 = blink present) with an equal number of samples as the pupil size data. Blink epochs were cut centered at stimulus and blank event onset times exactly as specified for the pupil size epochs (see *Pupil size epoch extraction* section), except that no baselining was performed on the blink epochs. Blink epochs were removed from analysis if more than 50 percent of the epoch samples between 1000 milliseconds pre-stimulus onset or blank event and 6000 milliseconds post-stimulus onset or blank were 0 (e.g., due to prolonged eye closure or loss of eye tracking). This epoch removal criterion resulted in the removal of less than 15 percent of blink epochs. Finally, epochs were averaged within participant across event type and stimulus presentation location and smoothed. The resulting mean blink fraction timecourses indicated the blink rate for each epoch sample.

#### Microsaccade epoch extraction

Microsaccade events were determined from the gaze position data acquired simultaneously with pupillometry (https://github.com/sj971/neurosci-saccade-detection; ^41^). The resulting microsaccade data set was a binary timecourse (0 = microsaccade absent; 1 = microsaccade present) with an equal number of samples as the pupil size data. Microsaccade epochs were cut centered at the stimulus onset and blank event times exactly as specified for the blink fraction epochs (see *Blink epoch extraction* section). Also, the same blink epoch removal criterion was applied to the microsaccade epochs resulting in rejecting less than 15 percent of microsaccade epochs. Finally, epochs were averaged within participant across event type and stimulus presentation location and smoothed. The resulting mean microsaccade fraction timecourses indicated the rate that a microsaccade event was present for each epoch sample.

#### Eye metric visualization and statistical analysis

For control participants, eye metric epoch – pupil size, blink, and microsaccade – timecourses were averaged between the left and right visual field stimulus presentation locations (Figure 3A; Supplementary Figure 3A). For patient participants, the eye metric epoch timecourses were averaged separately within the sighted and blind field stimulus presentation locations (Figure 3B, C, and D; Figure 5A; Supplementary Figure 3B, C, and D; Supplementary Figure 6). However, the blind field dynamics were visualized and statistically evaluated (see below) independently between blind aware and blind unaware patient participants (Figure 3C and D; Supplementary Figure 3C and D; Supplementary Figure 4C and D; see *Visual perception task* Visual behavior Results section for the definitions of blind aware and unaware).

**Figure 3.**
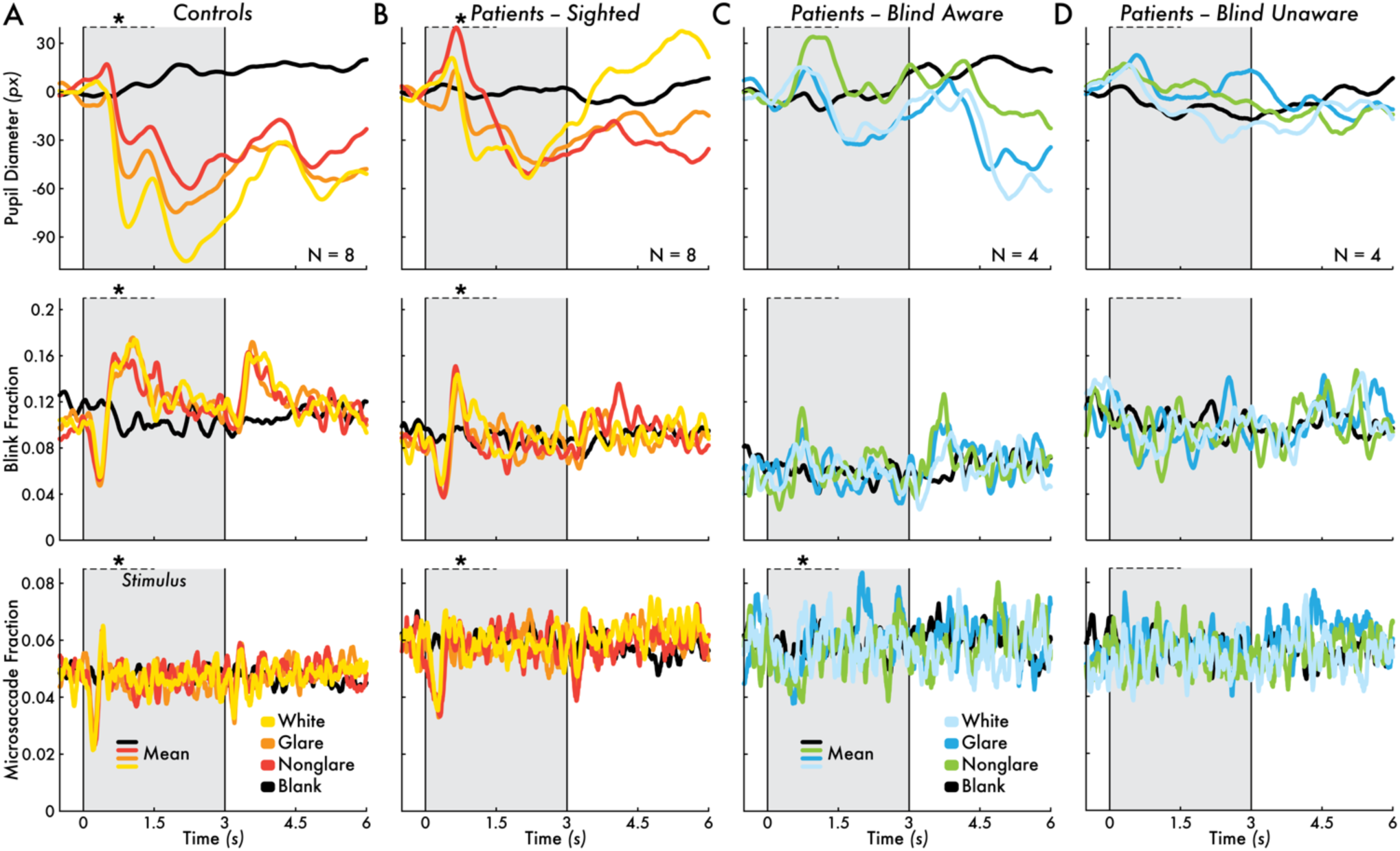
Nontarget stimulus-evoked pupil, blink, and microsaccade dynamics. Pupil diameter, blink fraction, and microsaccade fraction change preceding (0.5 seconds [s]) and following (6 s) the nontarget stimuli (white stimulus = yellow/light blue; glare stimulus = orange/blue; nonglare stimulus = red/green) or blank event (black) for **(A)** control participants (N = 8) averaged between the left and right visual field, **(B)** patient participants (N = 8) sighted field, **(C)** blind aware patient participants (N = 4) blind field, and **(D)** blind unaware patient participants (N = 4) blind field. In all subplots, the asterisks (*) indicates that there was a statistically significant (*p* < 0.05) difference in maximum or minimum eye metric signal change between the nontarget stimuli and blank event within the first 1.5 s from stimulus onset (horizontal dotted line; Supplementary Figure 4). The eye metric timecourses represent the group mean response. The 3-second stimulus presentation interval is highlighted by a gray area bounded between two vertical lines at 0 and 3 s.

Eye metric changes were statistically evaluated between the nontarget (glare, nonglare, and white stimulus) and blank event by finding the *minimum* and *maximum* eye metric value for each participant’s mean eye metric epoch timecourse within the first 1500 milliseconds after stimulus onset or blank event (the analysis interval represented by the horizontal dotted line in Figure 3; Supplementary Figure 4). A paired *t*-test (Holm-Bonferroni corrected for multiple comparisons) evaluated if the maximum and minimum pupil size values across the nontarget stimuli (mean response of glare, nonglare, and white stimulus) were statistically different from the corresponding blank events (mean response of blank glare, blank nonglare, and blank white events).

#### Eye metric-based classification

Epoch-level, stimulus-evoked eye metric dynamics were evaluated using participant-level classifiers trained on the pupil size, blink, and microsaccade epoch data. Four sets of classifiers were trained and tested for each participant. For control participants, the trained classifiers were: (1) left visual field and (2) right visual field *target stimulus* versus blank event, and (3) left visual field and (4) right visual field *nontarget stimulus* versus blank event. For patient participants, the trained classifiers were: (1) sighted field and (2) blind field *target stimulus* versus blank event, and (3) sighted field and (4) blind field *nontarget stimulus* versus blank event.

A two-step classification approach was implemented. First, pupil size, blink, and microsaccade linear support vector machine classifiers (i.e., three independent classifiers for each eye metric) were trained using 10-fold cross-validation on epoch samples within 4000 milliseconds post-stimulus onset. Next, the predicted scores (i.e., the signed distance from the decision boundary) for each epoch from the first-level pupil size, blink, and microsaccade classifiers were used as features in a second-level linear support vector machine classifier (i.e., a single classifier using the scores from all first-level classifiers as features). Finally, the classification results from the second-level classifier were used to assess classification performance. Both the first and second-level classifier were trained to predict epoch class: either a stimulus or blank event.

Classification performance was evaluated by *accuracy rate* (number of *correctly* predicted epochs divided by the number of *correctly* and *incorrectly* predicted epochs) and *area under the curve* (AUC) for receiver operating characteristics (ROC) curves (Figure 4; Supplementary Figure 5). Chance level accuracy rate was calculated for each participant and stimulus presentation location condition (chance level equals the total number of class 1 trials divided by all trials). In group-level analyses, a paired and one-sample *t*-test (Holm-Bonferroni corrected for multiple comparisons) evaluated if the accuracy rate and AUC, respectively, was statistically greater than chance (the AUC chance value was 0.5). Paired *t*-tests assessed if the accuracy rate and AUC performance were different between the left and right visual field (control participants) or the sighted versus blind field (patient participants).

**Figure 4.**
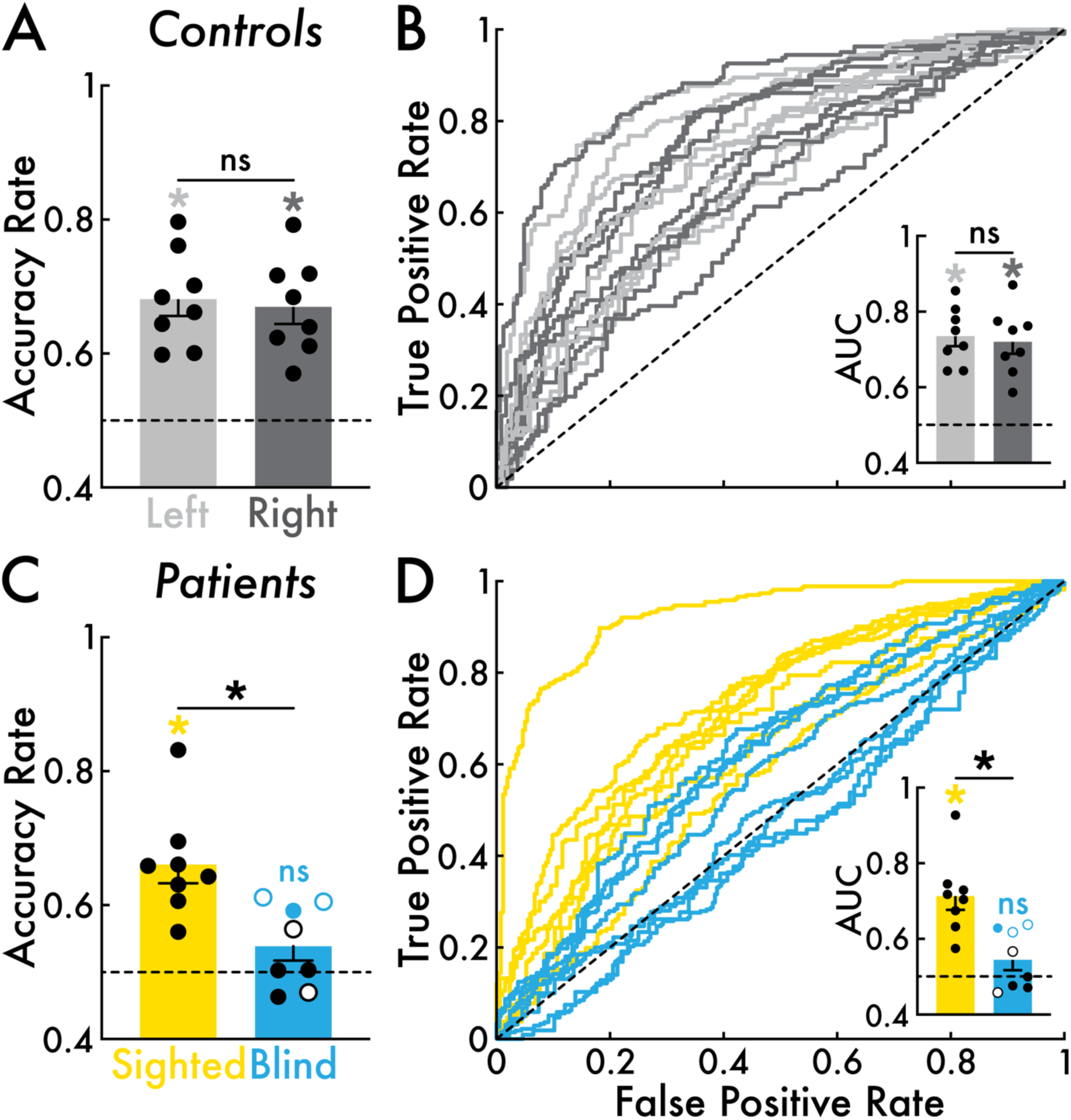
Eye metric-based nontarget stimulus versus blank event classification performance. **(A)** Control participants (N = 8) classification accuracy rate for the left (light gray) and right (dark gray) visual field. Left and right visual field accuracy rates were significantly (*; *p* < 0.05) greater than chance. Left versus right visual field accuracy rates were not significantly different (ns). **(B)** Control participants classification performance visualized by receiver operating characteristic (ROC) curves (false positive rate versus true positive rate). Each timecourse corresponds with a control participant and stimulus presentation location condition (left and right visual field). The inset bar graph depicts the area under the curve (AUC) for each ROC curve. Left and right visual field AUC were significantly (*p* < 0.05) greater than chance. Left versus right visual field AUC was not significantly different. **(C)** Patient participants (N = 8; blind aware patient participants N = 4 [open circles]; blind unaware patient participants N = 4 [closed circles]) classification accuracy rate for the sighted (yellow) and blind field (blue). Accuracy rate was significantly (*p* < 0.05) greater than chance for the sighted field but not significant for the blind field. Sighted field accuracy rate was significantly greater than the blind field (*p* < 0.05). Blind field accuracy rate was significantly (*p* < 0.05) greater than chance for three patient participants (highlighted with blue open and closed circles). **(D)** Patient participants classification performance visualized by ROC curves. Each timecourse corresponds with a patient participant and a location condition (sighted and blind field). The inset bar graph depicts the AUC for each ROC curve. AUC was significantly (*p* < 0.05) greater than chance for the sighted field but not significant for the blind field. Sighted field AUC was significantly (*p* < 0.05) greater than the blind field. Blind field AUC was significantly (*p* < 0.05) greater than chance for three patient participants (highlighted with blue open and closed circles). In all subplots, the open and closed circles represent individual participants, and the bars and error bars indicate the group mean and standard error of the mean, respectively. Chance level was 0.5 (highlighted with a dotted line).

Participant-level analyses were performed on the patient participants. Specifically, to evaluate whether accuracy rate and AUC were statistically greater than chance, a non-parametric percentile rank test was performed on the blind field target and nontarget classification performance (Figure 4C and D; Supplementary Figure 5C and D). First, a null distribution of accuracy rate and AUC was calculated by 500 random permutations of the epoch labels (i.e., stimulus or blank event epoch) and then running the eye metric classification procedure (see above) with the permuted labels. The percentile rank was calculated as the proportion of null distribution accuracy rate and AUC values *less than or equal* to the non-permuted epoch label accuracy rate and AUC performance.

#### Correspondence among blind field task behavior, verbal perceptual report, and eye metric dynamics

Three main measures were acquired from all patient participants: (1) behavioral performance on the visual perception task, (2) verbal report on perceptual experiences related to stimulus presentation during the visual perception task, and (3) stimulus-evoked eye metric dynamics. Evaluating the correspondence among these measures could help assess their interactions and robustness, particularly for indicating blind field visual conscious awareness. For patient participants, each measure type was evaluated as *present* or *absent* in the blind field. Task behavior was designated *present* if the target stimulus perception rate was > 0.25. Verbal report indicative of blind field conscious awareness was designated *present* if conscious vision, including degraded abnormal vision or non-visual sensations were reported for > 10 percent of task stimuli. Finally, stimulus-evoked eye metric dynamics was designated *present* in the blind field if stimulus evoked dynamics were observed by visual inspection in at least one of the eye metrics (pupil size, blink, or microsaccade; e.g., Figure 5A; Supplementary Figure 6) or above chance eye metric-based classification performance (e.g., blue open and closed circles in Figure 4C and D; see *Eye metric-based classification* section).

**Figure 5.**
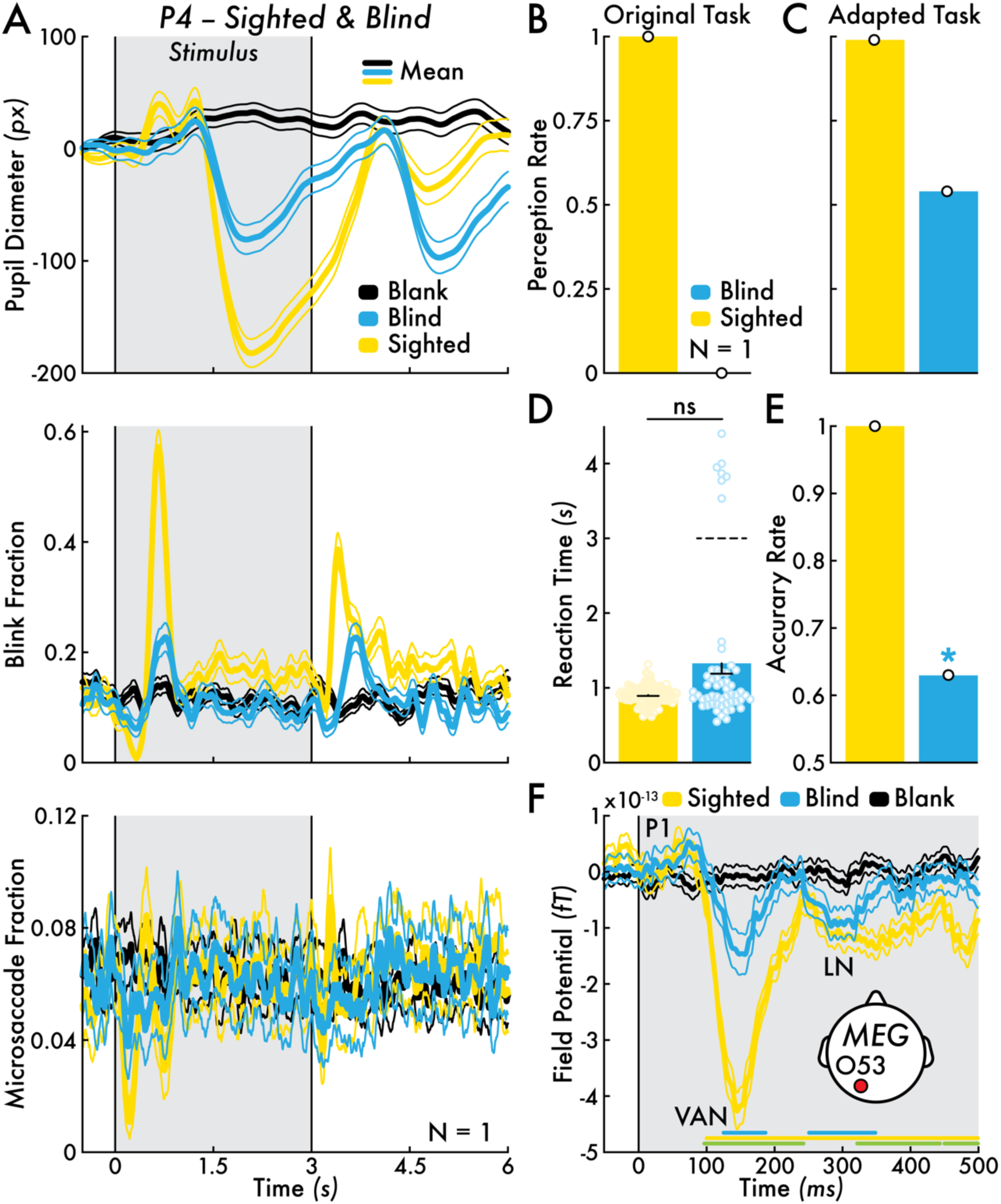
Patient participant P4 sighted and blind field behavior, eye metric dynamics, and MEG responses for nontarget stimulus. **(A)** Sighted (yellow) and blind field (blue) pupil diameter, blink fraction, and microsaccade fraction change for nontarget stimulus versus blank event (black). The mean eye metric changes across all nontarget stimulus trials are shown (thicker traces) bounded by the standard error of the mean (SEM; thinner traces). **(B)** Perception rate for the target stimuli in the sighted (yellow) and blind field (blue) for the *original* visual perception task (same result for P4 as in Figure 2B). **(C)** Perception rate for target stimuli in the sighted and blind field for the *behavioral-adapted* visual perception task (see *Visual Perception Task* Methods section). **(D)** Target stimulus response reaction time in the sighted (n = 99 responses) and blind (n = 54 responses) field for the behavioral-adapted visual perception task. The bar represents the mean reaction time with SEM. Individual trial reaction times are shown with open circles. A subset of responses (n = 7) may have corresponded with stimulus offset (horizontal dotted line). **(E)** Orientation discrimination accuracy rate for target stimuli in the sighted and blind field for the behavioral-adapted visual perception task. The blind field accuracy rate was significantly (*; *p* < 0.05) above chance. **(F)** Nontarget stimuli and blank events evoked magnetencephalography (MEG) field potentials (femtotesla; fT) in the sighted (yellow) and blind field (blue) for representative left occipital sensor O53 (see inset diagram for approximate sensor location on the scalp; see Supplementary Figure 7 for results from all left and right occipital sensors). Major field potential components are highlighted: first positivity (P1), N2 or visual awareness negativity (VAN), and late negativity (LN). Statistically significant (*p* < 0.05) intervals determined by cluster-based permutation testing are shown with horizontal lines for three evaluated contrasts: (1) sighted nontarget stimulus versus sighted + blind field blank events (yellow), (2) blind nontarget stimulus versus sighted + blind field blank events (blue), and (3) sighted versus blind field nontarget stimulus (green). The mean field potential changes across all nontarget stimulus trials are shown (thicker traces) bounded by SEM (thinner traces). The MEG results were acquired with the *MEG-adapted* visual perception task (see *Visual Perception Task* Methods section). Note that no error bars are depicted in subplots B, C, and D because the bars represent a single participant value: P4’s mean perception rate and accuracy rate.

### MEG

MEG analyses were completed using custom functions and the FieldTrip toolbox (http://fieldtriptoolbox.org; ^42^). First, the MEG data were preprocessed on a sensor basis, including applying a bandpass filter (0.1 and 115 Hz) and line noise filter (60 and 120 Hz). Next, the preprocessed MEG data were cut into 8001-millisecond epochs centered at stimulus onset and blank event times (see *Visual Perception Task* section). Stimulus event times were determined by analyzing the parallel port triggers that were recorded simultaneously with the MEG sensors. An event time correction of 19 milliseconds was applied to the trigger onset times to correct for the delay between the parallel port trigger time and the presentation of task stimuli on the projector screen (see *Testing Equipment and Facility* section). The blank event times were determined using the same method specified for pupil size blank event epochs (see *Pupil epoch extraction* section). Finally, sensor-level field potential averages and standard error of the mean across epochs were calculated within event type and sighted and blind field stimulus presentation locations. The MEG results reported in Figure 5F and Supplementary Figure 7 represent the average field potential across all nontarget stimuli (glare, nonglare, and white stimulus) in the sighted (yellow) and blind field (blue), while the blank event timecourse (black) represents an average between the sighted and blind field.

Sensor-level statistically significant changes in field potential among the sighted nontarget, blind nontarget, and sighted + blind field blank event conditions were performed with cluster-based permutation testing (5000 permutations; ^43^). The permutation analysis baseline interval was 500 milliseconds preceding the stimulus onset or blank event. The 500 milliseconds following the stimulus onset or blank event were evaluated for statistically significant samples. Three statistical comparisons were made: (1) sighted nontarget stimuli versus sighted + blind field blank events, (2) blind nontarget stimuli versus sighted + blind field blank events, and (3) sighted nontarget stimuli versus blind field nontarget stimuli (Figure 5F). Statistical testing was not performed on the sensors depicted in Supplementary Figure 7.

## Results

### Demographics

Control and patient participants did not statistically differ in age or education (*p* > 0.05). This result supports that the control and patient participants were matched by age and education.

### Visual behavior

#### Visual perception task

Control participants performed with high target stimulus perception rate (left visual field mean rate = 0.96; right visual field mean rate = 0.95; Figure 2A) and high target stimulus orientation discrimination accuracy rate (left visual field mean rate = 0.95; right visual field mean rate = 0.97; Figure 2C). Left versus right visual field perception rate and accuracy rate did not statistically differ (*p* > 0.05).

For stimuli presented in the sighted field, patient participants performed with high target stimulus perception rate (mean rate = 0.95; Figure 2B) and high target stimulus orientation discrimination accuracy rate (mean rate = 0.86; Figure 2D). However, for target stimuli presented in the blind field, the patient participants performed with low perception rate (mean rate = 0.26; Figure 2B), except for patient participants P2 and P7 with perception rates greater than 0.9. Excluding P2 and P7, the group mean perception rate for target stimuli in the blind field was 0.043. The blind field perception rate was significantly less than the sighted field (*t*[7] = 4.59, *p* < 0.0025). The blind field target stimulus orientation discrimination accuracy rate was 0.88 and 0.57 for P2 and P7, respectively (Figure 2D).

All participants reported instances of either residual or degraded conscious vision or non-visual sensations related to stimulus presentation, including reports of seeing “faint gray” (P1), “movement” (P4), and “very slight shadows” (P5), and non-visual sensations like “felt an image” (P6; Table 3 and 4). Patient participants were categorized according to their verbal report as either (1) *blind aware* or (2) *blind unaware*. The blind *aware* patient participants (P2, P4, P7, and P8) frequently (> 10 percent of trials) reported conscious awareness in the blind field, including the statements “barely could see images” (P2), “I could sense there was movement” (P4), “confident about color but not shape” (P7), and “barely only able to catch the left [blind field] images” (P8; Table 3). The blind *unaware* patient participants (P1, P3, P5, and P6) infrequently (< 10 percent of trials) reported conscious awareness in the blind field (Table 4). In some cases, conscious awareness for stimuli in the blind field for blind unaware patient participants were related to eye movements (i.e., making saccades away from central fixation).

**Table 4.** Summary of *blind unaware* patient participants’ blind field task behavior, eye metrics, and verbal perceptual report results (see *Correspondence among blind field task behavior, verbal perceptual report, and eye metric dynamics* Methods section). Note that verbal report was considered absent when visual conscious awareness was reported in < 10 percent of trials. Some instances of blind field conscious awareness were linked to eye movements away from central fixation.

| Blind Unaware |  |  |  |  |
| --- | --- | --- | --- | --- |
| *Patients* | *Task Behavior* | *Eye Metrics* | *Verbal Report* | *Verbal Report Summary* |
| *P1* | Absent | Absent | Absent | <u>Summary:</u> ~10 instances of seeing or sensing task stimuli.<br><u>Representative quotes:</u> “might have felt [an image] but so faint and fast”; “saw a faint gray [image]”; “might have noticed a red or gray image [but might be] my mind playing tricks on me”; “might have seen silver images on the left [blind field] but just might have been my brain”. |
| *P3* | Absent | Absent | Absent | <u>Summary:</u> ~6 instances of seeing or sensing task stimuli.<br><u>Representative quotes:</u> “I did not see anything to my right [blind field] but had the feeling to look to the right as if something might have appeared”; “an image on the right [blind field] just appeared in my view”; “it [the image] kind of popped up”. |
| *P5* | Absent | Present | Absent | <u>Summary:</u> ~12 instance of seeing task stimuli.<br><u>Representative quotes:</u> saw “very slight shadows”, “white shadow”, “red tint”, and “red shadow on the left [blind field]”; “could see red [in the blind field] but not sure what it was”; “saw red [in the blind field] but could not tell the shape”. |
| *P6* | Absent | Absent | Absent | <u>Summary:</u> ~9 instance of seeing or sensing task stimuli.<br><u>Representative quotes:</u> “felt once that there was something on the right [blind field]”; “felt an image [in the blind field] but not sure; was not vision”. |

Patient participant P4 performed with a perception rate of 1 and 0 for target stimuli in the sighted and blind field, respectively (Figure 2B; Figure 5B). However, the presence of blind field, stimulus-evoked eye metric responses (see *Eye metrics* section; Figure 5A) prompted a subsequent interview with P4 on his blind field visual conscious experiences. P4 described that he occasionally saw or felt “movement”, “clicking down”, “vibration”, or the addition of static noise in his blind field (Table 3). These experiences were distinct from normal conscious vision. As P4 explained, “I could sense there was movement in the upper right [blind field] but could not see anything”. Notably, P4 believed his blind field conscious experiences occurred at random, so did not correspond with visual sensory stimulation.

A behavioral-adapted visual perception task was developed to probe whether P4’s blind field experiences corresponded with stimulus presentation (see *Visual Perception Task* Methods section). In the adapted task, P4 was instructed to make keypresses whenever he saw a target stimulus or experienced a sensation in his blind field (e.g., the feeling of movement). P4’s blind field perception rate on the new behavioral-adapted visual perception task was 0.54 – an increase from a perception rate of 0 in the original visual perception task (Figure 5B versus C). Notably, P4’s blind field false positive rate (i.e., reporting a stimulus when none was present) was 0. Moreover, P4’s sighted versus blind field reaction time (RT) relative to stimulus onset were not statistically different (sighted field mean RT = 0.91 seconds; blind field mean RT = 1.33 seconds; *p* > 0.05; Figure 5D). There was a subset of trials (n = 7) with prolonged RT, clustered at approximately 4 seconds post-stimulus onset. One plausible explanation is that these late responses corresponded with stimulus *offset.* These results indicated that P4’s blind field conscious experiences directly corresponded with the presentation of task stimuli.

P4’s target stimulus orientation discrimination performance was poor (34 correct responses out of 54 responses; accuracy rate = 0.63), although significantly above chance (*p* < 0.038; Figure 5E). P4 indicated that he guessed the stimulus orientation by “instinct”, “inspired guess”, and by “an electric charge coming down my finger” so that even “before I could think, I pressed the button”. P4’s descriptions for how he determined the stimulus orientation in his blind field are similar to reports by people with cerebral blindness who also achieve above chance performance on visually guided tasks by “only guessing” ^4^. Overall, P4’s blind field behavioral performance corresponds with stimulus-evoked eye metric responses and MEG field potentials (see *Eye metrics* and *Blind field residual brain processing* sections; Figure 5A and F; Supplementary Figure 7).

#### Brightness perception task

Control and patient participants reported similar perceived brightness of the glare, nonglare, and isoluminant stimulus. A one-way ANOVA revealed a statistically significant effect of stimulus type on perceived brightness for control (F[1.02, 7.13] = 149.5, *p* < 0.0001) and patient participants (F[1.13, 7.89] = 26.62, *p* = 0.0007). Post-hoc paired *t*-test analyses indicated that the glare stimulus was reported as significantly brighter than the nonglare stimulus for control participants (*t*[7] = 5.07, *p* = 0.0015; patient participants *p* > 0.05), and the nonglare stimulus was reported as significantly brighter than the isoluminant stimulus for control (*t*[7] = 15.28, *p* < 0.0001) and patient participants (*t*[7] = 8.67, *p* < 0.0001; Supplementary Figure 2B and C).

### Eye metrics

#### Target stimuli

Control participants showed robust changes in pupil size, blink rate, and microsaccade rate for target stimuli versus corresponding blank events. On the group-level, target stimuli induced a unimodal pupil dilation after stimulus onset, blink rate suppression after stimulus onset and blink rate enhancement after stimulus offset, and microsaccade rate suppression after stimulus onset and offset (Supplementary Figure 3A).

The group-level eye metric dynamics corresponded with high, statistically above chance epoch-level classification performance for discriminating target stimulus versus blank event epochs based on the pupil size, blink rate, and microsaccade rate responses for both the left (mean accuracy rate = 0.80; *t*[7] = 30.32, *p* < 0.0001) and right visual field (mean accuracy rate = 0.83; *t*[7] = 12.08, *p* < 0.0001; Supplementary Figure 5A). Also, ROC AUC performance was statistically greater than chance for the left (mean AUC = 0.86; *t*[7] = 8.74, *p* < 0.0001) and right visual field (mean AUC = 0.89; *t*[7] = 7.03, *p* = 0.0002; Supplementary Figure 5B). The left versus right visual field accuracy rate and ROC AUC performance did not statistically differ (*p* > 0.05). These results support that the target stimulus evoked eye metric dynamics are similar between the left and right visual field stimulus presentation locations.

Patient participants revealed similar pupil size, blink rate, and microsaccade rate changes as control participants for target stimuli presented in the sighted field (Supplementary Figure 3A versus B). However, there were differences observed between the control and patient participant sighted field versus the patient participant blind field eye metrics dynamics. For blind aware patient participants, the blind field eye metric dynamics were reduced in amplitude and response duration yet shared a similar profile (e.g., pupil dilation and microsaccade rate suppression following stimulus onset; Supplementary Figure 3C). For blind unaware patient participants, there were no changes in pupil size, blink rate, or microsaccade rate (Supplementary Figure 3D).

The group-mean eye metric timecourses corresponded with high and low classification performance for discriminating target stimulus versus blank event epochs based on the eye metric responses in the sighted and blind fields, respectively (sighted field mean accuracy rate: 0.81; blind field mean accuracy rate: 0.61; Supplementary Figure 5C and D). Classification accuracy rate and ROC AUC were significantly above chance in the sighted field (accuracy rate: *t*[7] = 15.77, *p* < 0.0001; AUC: *t*[7] = 18.37, *p* < 0.0001). Classification performance was also significantly different from chance in the blind field (accuracy rate: *t*[7] = 2.59, *p* = 0.036; AUC: *t*[7] = 2.62, *p* = 0.035), largely driven by the blind aware patient participants (see details below). The sighted versus blind field accuracy rate and AUC performance statistically differed (accuracy rate: *t*[7] = 4.78, *p* = 0.002; AUC: *t*[7] = 4.04, *p* = 0.0049; Supplementary Figure 5C and D).

Participant-level analyses of the blind field classification performance revealed that four of four blind aware (P2, P4, P7, and P8) and one of four blind unaware patient participants (P5) achieved statistically significant (*p* < 0.05) above chance accuracy rate and AUC (except P8 who was not above chance for AUC performance; blue open [blind aware] and closed circles [blind unaware] in Supplementary Figure 5C and D).

#### Nontarget stimuli

Control participants showed robust changes in pupil size, blink rate, and microsaccade rate for nontarget stimuli (glare, nonglare, and white stimulus) versus corresponding blank events. Specifically, nontarget stimuli induced sustained pupil constriction after stimulus onset, blink rate suppression and enhancement after stimulus onset and blink rate enhancement after stimulus offset, and microsaccade rate suppression and enhancement after stimulus onset. Pupil size change was modulated by the nontarget stimulus type. Specifically, the largest and smallest constriction was observed for the white and nonglare stimuli, respectively. However, the blink and microsaccade fraction change was not modulated by stimulus type. Nontarget stimulus versus blank event maximum signal change was statistically significant for blink (*t*[7] = 3.75, *p* = 0.0072) and microsaccade rate (*t*[7] = -2.80, *p* = 0.027); minimum signal change was statistically significant for pupil size (*t*[7] = 2.57, *p* = 0.037) and microsaccade rate (*t*[7] = 3.96, *p* = 0.0055; Figure 3A; Supplementary Figure 4A).

The group-level eye metric dynamics corresponded with moderate, statistically above chance epoch-level classification performance for discriminating nontarget stimulus versus blank event epochs based on the pupil size, blink rate, and microsaccade rate responses for both the left (mean accuracy rate = 0.68; *t*[7] = 7.166, *p* = 0.0002) and right visual fields (mean accuracy rate: 0.67; *t*[7] = 6.85, *p* = 0.0002; Figure 4A). Also, ROC AUC performance was statistically greater than chance for the left (mean AUC = 0.74; *t*[7] = 8.74, *p* < 0.0001) and right visual field (mean AUC = 0.72; *t*[7] = 7.03, *p* = 0.0002; Figure 4B). The left versus right visual field accuracy rate and AUC performance did not statistically differ (*p* > 0.05). These results support that the nontarget stimulus-evoked eye metric dynamics are similar between the left and right visual field stimulus presentation locations.

Patient participants revealed similar pupil size, blink rate, and microsaccade rate changes as control participants for nontarget stimuli presented in the sighted field (Figure 3A versus B; Supplementary Figure 4A versus B). Eye metric responses included pupil constriction after stimulus onset, blink rate suppression and enhancement after stimulus onset and blink rate enhancement after stimulus offset, and microsaccade rate transient suppression and enhancement after stimulus onset (Figure 3B). As with the control participants, pupil size but not blink and microsaccade rate change was modulated by nontarget stimulus type, with the greatest and smallest pupil constriction observed for the white and nonglare stimulus, respectively. Sighted field nontarget stimulus versus blank event minimum signal change was statistically significant for pupil size (*t*[7] = 3.055, *p* = 0.018), blink rate (*t*[7] = 3.41, *p* = 0.011) and microsaccade rate (*t*[7] = 3.38, *p* = 0.012; Figure 3B; Supplementary Figure 4B). Pupil size, blink rate, and microsaccade rate maximum signal change were not statistically significant (*p* > 0.05)

However, there were differences observed between the control and patient participant sighted field versus the patient participant blind field eye metrics dynamics. For blind aware patient participants, the blind field eye metric dynamics were reduced in amplitude and response duration yet shared a similar profile (e.g., pupil constriction, blink rate enhancement, and microsaccade rate suppression following stimulus onset; Figure 3C). However, only the minimum microsaccade rate change was statistically significant for the blind field nontarget stimulus versus blank event (*t*[3] = 4.49, *p* = 0.021; Figure 3C; Supplementary Figure 4C). Notably, visual inspection showed that the blind aware pupil size change no longer discriminated between the white and glare stimulus. For blind unaware patient participants, there were no changes in pupil size, blink rate, or microsaccade rate (Figure 3D; Supplementary Figure 4D).

The sighted and blind field eye metric dynamics corresponded with eye metric-based, epoch-level classification performance of nontarget stimulus versus blank event. Statistically above chance classification accuracy rate (mean accuracy rate = 0.66; *t*[7] = 5.85, *p* = 0.0006; Figure 4C) and AUC (mean AUC = 0.71; *t*[7] = 5.85, *p* = 0.0006; Figure 4D) was found for discriminating between nontarget stimuli and blank event epochs in the sighted field. The sighted versus blind field accuracy rate and AUC performance statistically differed (accuracy rate: *t*[7] = 4.16, *p* = 0.0042; AUC: *t*[7] = 4.21, *p* = 0.004).

Participant-level analyses of the blind field classification performance revealed that two of four blind aware (P2 and P4) and one of four blind unaware patient participants (P5) achieved statistically significant (*p* < 0.05) above chance accuracy rate and AUC (blue open [blind aware] and closed circles [blind unaware] in Figure 4C and D).

### Blind field visual cortical processing

Visual stimulus-evoked MEG field potentials were recorded for patient participant P4. Scalp field potential changes over the left occipital cortex for nontarget stimuli presented in the sighted field included a small increase at 100 milliseconds (first positivity or P1), a large negative change between 100-225 milliseconds (N2 or visual awareness negativity [VAN]), and subsequent late negative (LN) changes from 250 milliseconds (Figure 5F; Supplementary Figure 7A). Smaller amplitude but similarly timed field potential changes were also found for the nontarget stimuli presented in the blind field. Scalp sensors over the right occipital cortex revealed weaker, but similarly timed responses, predominantly for stimuli presented in the sighted field (Supplementary Figure 7B).

Cluster-based permutation testing on representative left occipital sensor O53 (likely responsive to neural activity in the left visual cortex; see Figure 5F for approximate sensor location on the scalp) highlighted that the stimulus-evoked changes were greater than blank event for both the sighted and blind field. However, the field potential response amplitude was significantly greater in the sighted versus blind field, particularly during the VAN and LN intervals. It is also notable that the sighted and blind field potential profile matched the sighted versus blind field eye metrics dynamics (Figure 5A versus F). Two compatible explanations for the amplitude difference in the sighted and blind eye metric and MEG responses are: (1) the averaged responses combine perceived and not perceived stimuli (i.e., P4 reported stimulus-evoked conscious experiences in his blind field for ∼50 percent of trials; Figure 5C), and (2) impaired visual neural processing.

#### Correspondence among blind field task behavior, verbal perceptual report, and eye metric dynamics

Task behavior, verbal report, and eye metrics in the blind field agreed in six patient participants: all present (blind aware: P2, P4, and P7; Table 3) and all absent (blind unaware: P1, P3, and P6; Table 4). There was disagreement among the measures for blind aware P8 and blind unaware P5. For P8, a pupillary response and verbal report were present (Supplementary Figure 6B; Table 3), while task behavior indicative of blind field conscious awareness was absent (blind field perception rate = 0.06). However, P8’s verbal reports indicated that his poor behavioral performance was related to impaired color vision in his blind field (e.g., “what might be red is grayish”) making it difficult to discriminate between target and non-target stimuli (Table 3). However, P8’s blind field eye metric classification performance was significantly (*p* < 0.05) above chance for only target accuracy rate (Supplementary Figure 5C).

For P5, a post-stimulus onset blink and microsaccade rate suppression were present in the blind field that matched the amplitude and timing of the sighted field blink and microsaccade dynamics (Supplementary Figure 6A). These eye metric responses, apparent by visual inspection, corresponded with significantly (*p* < 0.05) above chance eye metric classification performance (target accuracy rate = 0.65; target AUC = 0.65; nontarget accuracy rate = 0.59; nontarget AUC = 0.63; blue closed circles in Figure 4C and D and Supplementary Figure 5C and D). Nevertheless, P5 is categorized as blind unaware because she rarely verbally reported visual conscious awareness in her blind field, including no instances of non-visual sensations of the task stimuli (e.g., “very slight shadows”; Table 4). If this categorization is accurate, P5’s blind field eye metric responses may be linked to unconscious, residual neural processing of the task stimuli.

## Discussion

We evaluated visual stimulus-evoked pupil size, blink, and microsaccade dynamics in healthy and cerebrally blind participants. A key finding was that eye metric responses were linked with visual conscious awareness in both the sighted and blind field. Notably, eye metrics were present independent of behavioral performance on a visual perception task. Thereby, eye metrics may predict blind field conscious awareness even when behavior and verbal report suggest blindness.

We also observed individual differences in the presence of blind field eye metrics. For instance, patient participant P5 retained blink and microsaccade responses in their blind field (Supplementary Figure 6A); P8 retained only pupil responses (Supplementary Figure 6B); and P4 retained pupil, blink, and microsaccade responses (Figure 5A). This finding emphasizes the value of recording *multiple* eye metrics in the same person due to heterogenous responsiveness. Furthermore, a novel methodological outcome from this study is to show that machine learning methods can provide a robust, objective approach for detecting stimulus-level eye metric responses.

The maintenance of stimulus-evoked eye metric dynamics in the blind field suggests residual neural processing of visual stimuli. In support, we found similar patterned MEG potential changes for visual stimuli presented in the sighted and blind field, including the field components P1, N2 or VAN, and LN (Figure 5F; Supplementary Figure 7A) ^44^. The VAN has been linked with visual conscious perception, while earlier and later brain potentials may be related to unconscious and post-perceptual processing ^22,45^.

In addition, we replicated a previous finding that perceived image brightness is linked to pupil size constriction (Figure 3A and B; Supplementary Figure 4A and B). Our results offer the additional insight that pupil size modulation based on perceived brightness depends on conscious perception. Also, blinking and microsaccade change did not discriminate perceived brightness. For blind aware patient participants, eye metrics changes were presented in the blind field, including pupillary responses that distinguished between perceived bright and less bright stimuli. Meanwhile, on the group level, there were no eye metric changes in the blind field for blind unaware patient participants. However, blind unaware patient participant P5 did reveal blind field stimulus-evoked blink and microsaccade suppression similar to her sighted field responses, which corresponded with above chance eye metric-based classification performance for target and nontarget stimuli (Figure 4C and D; Supplementary Figure 5C and D; Supplementary Figure 6A). This may be a case where stimulus-evoked eye metric dynamics and conscious awareness were decoupled (see *Limitations* section).

The absence of a pupillary light response to stimuli presented in the blind field of blind unaware patient participants agrees with previous reports of impaired pupillary light response in cerebral blindness ^34^. Another key factor is the intensity of light stimulation required to induce the pupillary light response. A previous report found some people with cerebral blindness only demonstrated a pupillary light response after dark adaptation and to very bright light stimulation (e.g., direct sunlight) ^33^. Therefore, the experimental parameters of the current study may have been insufficient to evoke a blind field pupillary light response.

### Blind field conscious awareness in cerebral blindness

A major source of controversy in the study of cerebral blindness is what exactly do these people experience in their blind field. Interrogating blind field conscious awareness is challenging because residual and degraded conscious vision or non-visual sensations may be reported as *no* visual conscious awareness. Therefore, the method for inquiring on perceptual experiences in cerebral blindness is influential ^19,46^. For instance, graded perceptual scales may be more sensitive than binary questionnaires (e.g., did you *see* something or not) ^13,47^.

Our results highlight this concern because patient participant P4 reported no conscious vision in the original visual perception task. However, robust eye metric responses to stimuli presented in his blind field (Figure 5A) prompted a subsequent interview and testing an adapted visual perception task, both revealing blind field conscious awareness for task stimuli that he reported as a “feeling of movement” or “vibration” (Figure 5B versus C; Table 3). P4 was also above chance in blind field stimulus orientation discrimination, although he decided by “instinct” or “inspired guess” (Figure 5E). Notably, P4 was unaware these blind field conscious experiences were indicative of visual sensory stimulation. This shares a new challenge for the method of subjective report to gauge conscious awareness in cerebrally blind people. Like P4, some patients may be unaware that their blind field conscious awareness is related to vision. Correspondingly, these patients may report that they are blind because their blind field perceptual experiences are not recognized as linked to visual stimulation.

Furthermore, patient participant P8’s performance on the visual perception task indicated little to no conscious vision in the blind field. However, P8 revealed by verbal report that he frequently perceived task stimuli in his blind field but was unable to distinguish the red target versus achromatic, nontarget stimuli (Table 3). Therefore, he decided not to respond to any stimulus during the task. Even without P8’s verbal report, blind field conscious awareness could be inferred by a robust stimulus-evoked pupillary response that was similar to the sighted field (Supplementary Figure 6B). P4 and P8 highlight that relying on behavioral performance or verbal report alone can neglect cases of blind field conscious awareness.

### Limitations

While our patient group is diverse by age, education, and ethnicity, the current findings may not generalize to other people with cerebral blindness due to unique injury profile (e.g., lesion severity, duration, and location). Also, heterogenous responses according to task and stimulus type is a major challenge for establishing consensus in cerebral blindness research. Stimulus features (e.g., size, duration, spatial frequency, luminance, color, and contour) can influence conscious awareness and eye metrics in cerebral blindness ^4,12,33,48,49^. Likewise, the current stimulus parameters may evoke unique behavior, eye metrics, and neural responses that are not replicated with other stimulus types.

Furthermore, a limitation of eye metrics is that they are not conclusive of conscious awareness. For instance, some people with cerebral blindness show behavioral evidence of blindsight without corresponding pupillary responses ^35^. Likewise, we found that eye metrics were not in agreement with task behavior and verbal report in the blind unaware patient participant P5: task behavior and verbal report indicated no blind field conscious awareness, while eye metric responses were present (Figure 4C and D; Supplementary Figure 5C and D; Supplementary Figure 6A; Table 4). Accordingly, eye metrics should be considered *one* source of evidence among other data points, including behavior and neuroimaging in the assessment of conscious awareness. Likewise, the presence of neural activity patterns linked with conscious perception (e.g., the VAN; Figure 5F) may help guide assessment of conscious awareness in cerebral blindness.

## Conclusions

Probing visual conscious awareness and residual neural processing in cerebral blindness is a long-standing query. In the current study, pupil size, blink, and microsaccade dynamics are shown to be an accessible, objective measure of visual conscious awareness and brain processing in cerebral blindness. The results also highlight previous concerns that behavioral performance on a visual task may neglect blind field conscious awareness, particularly those characterized as degraded abnormal vision and non-visual sensations. We also show that each participant has a unique combination of eye metrics, thereby recording multiple measures is recommended. Future translational applications of eye metrics in cerebral blindness includes identifying opportunities for rehabilitation and tracking recovery of blind field conscious awareness and residual neural processing.

## Code and Data Availability

Data and code are available at https://github.com/nimh-sfim/eye_metrics_cerebral_blindness (will be made public upon publication).

## Author Contributions

S.I.K.: conceptualization, methodology, software, validation, formal analysis, investigation, data curation, writing (original draft), visualization, supervision, and project administration; V.E.G.: methodology, software, investigation, resources, writing (original draft), and visualization; S.J.: methodology, writing (review & editing), and project administration; E.M.: conceptualization, methodology, and writing (review & editing), funding acquisition; B.O.: resources and writing (review & editing); P.A.B.: methodology, funding acquisition, writing (review and editing), and supervision; T.T.L.: conceptualization, methodology, resources, writing (review & editing), supervision, and project administration.

## Acknowledgements

This research was supported by the Intramural Research Program of the National Institute of Mental Health (ZIAMH002783 to P.A.B. and ZIAMH002966 to E.P.M.). The study was completed in compliance with the National Institutes of Health Clinical Center protocol ID 10-M-0047 (ClinicalTrials.gov ID: NCT01087281). We thank the patients and their families for their time and participation in the study, Francisco Pereira for his insights on the machine learning procedures, Jeff Stout for his guidance on the MEG preprocessing and visualization procedures, and Mike Reel, Brittany Pollard, and the OP4 Behavioral Health Clinic staff for their support with patient admission and physical exams.

---

**Supplementary Figure 1.**
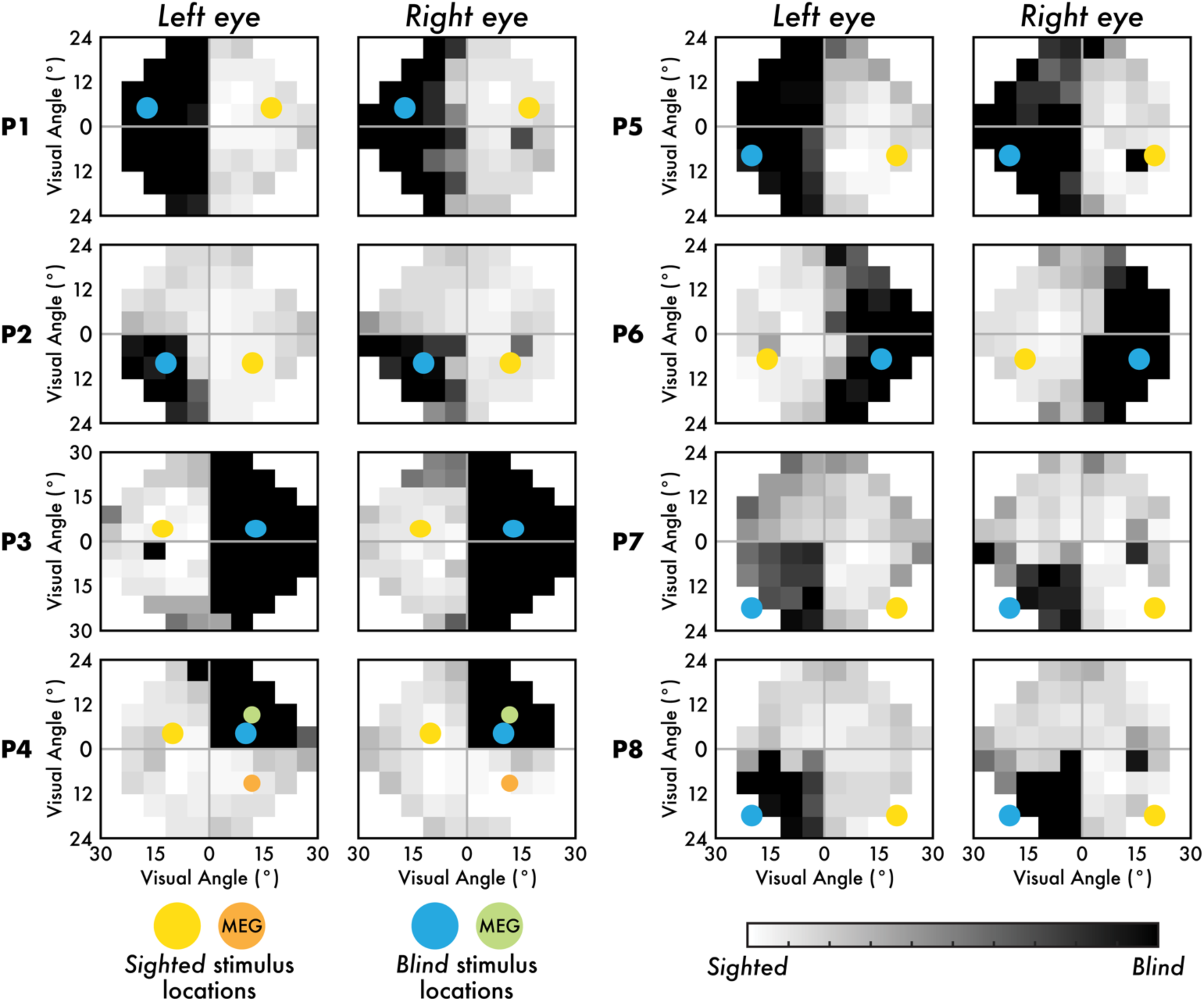
Humphrey visual field test results and sighted and blind field stimulus presentation locations. The Humphrey visual field (HVF) test results are shown for the left and right eye in degrees (°) of visual angle. Sighted areas are indicated by white and light gray squares, while blind areas are indicated by dark gray and black squares. The single black or dark gray square present for most patient participants in the left visual field (left eye) or the right visual field (right eye) is the natural blind spot. The colored circles depict the approximate size and location of the visual perception task stimuli shown in the sighted (yellow; MEG study session: orange) and blind field (blue; MEG study session: green).

**Supplementary Figure 2.**
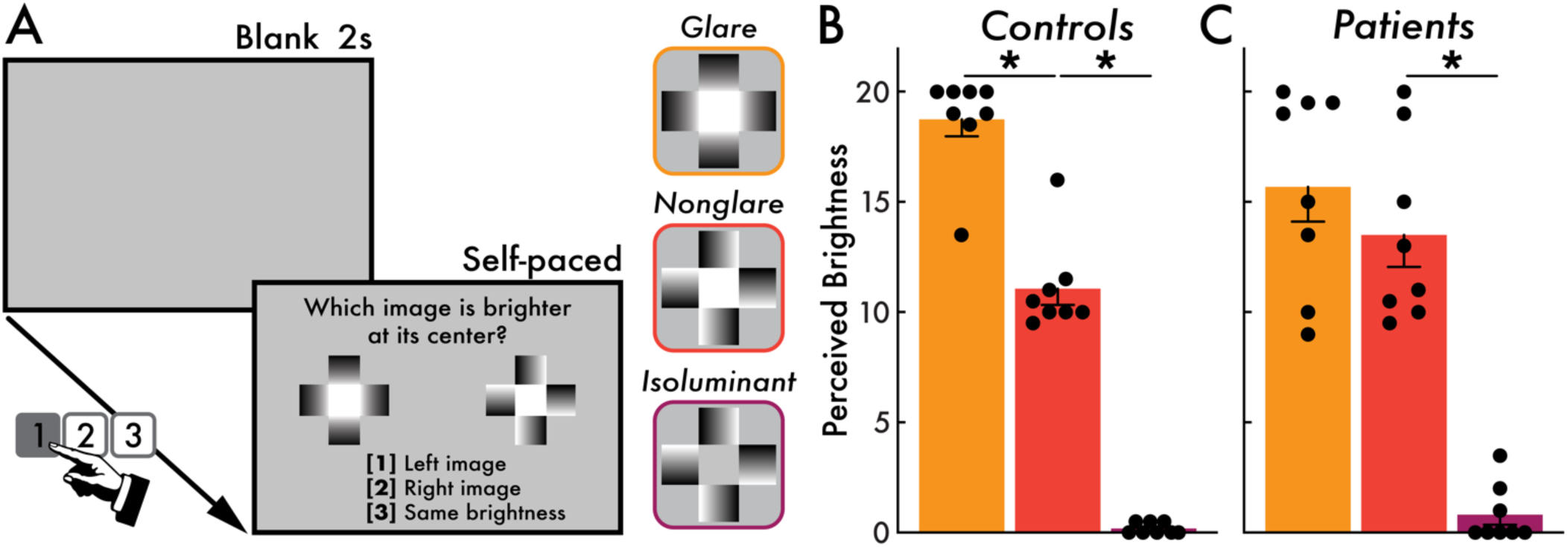
Brightness perception task and behavioral results. **(A)** Brightness perception task trial structure. First, a 2-second (s) blank interval appeared. Next, participants were shown two stimuli among three possible stimulus types: (1) glare, (2) nonglare, and (3) isoluminant. Participants were prompted to answer the question “Which image is brighter at its center?” Participants could view these stimuli directly and for an unlimited duration. Participants selected the 1-key if the left image was perceived as brighter, the 2-key if the right image was perceived as brighter, and the 3-key if both images appeared with equal brightness. Once the participant made their response, a new trial would begin with an updated image comparison set. Participants completed 30 trials total with 10 trials each of the following stimulus comparisons: (1) glare versus nonglare, (2) glare versus isoluminant, and (3) nonglare versus isoluminant. **(B)** Control participants (N = 8) perceived brightness for the glare (orange), nonglare (red), and isoluminant (purple) stimulus. Larger values (maximum = 20; minimum = 0) indicated that participants perceived that stimulus as brighter. The glare stimulus was reported as significantly brighter than the nonglare stimulus, and the nonglare stimulus was reported as significantly brighter than the isoluminant stimulus (*; *p* < 0.05). **(C)** Patient participants (N = 8) perceived brightness for the glare, nonglare, and isoluminant stimulus. Most patient participants reported that the glare stimulus was brighter than the nonglare stimulus, however, this response was not statistically significant. The nonglare stimulus was reported as significantly (*p* < 0.05) brighter than the isoluminant stimulus. In B and C, the circles represent individual participants, and the bars and error bars indicate the group mean and standard error of the mean, respectively.

**Supplementary Figure 3.**
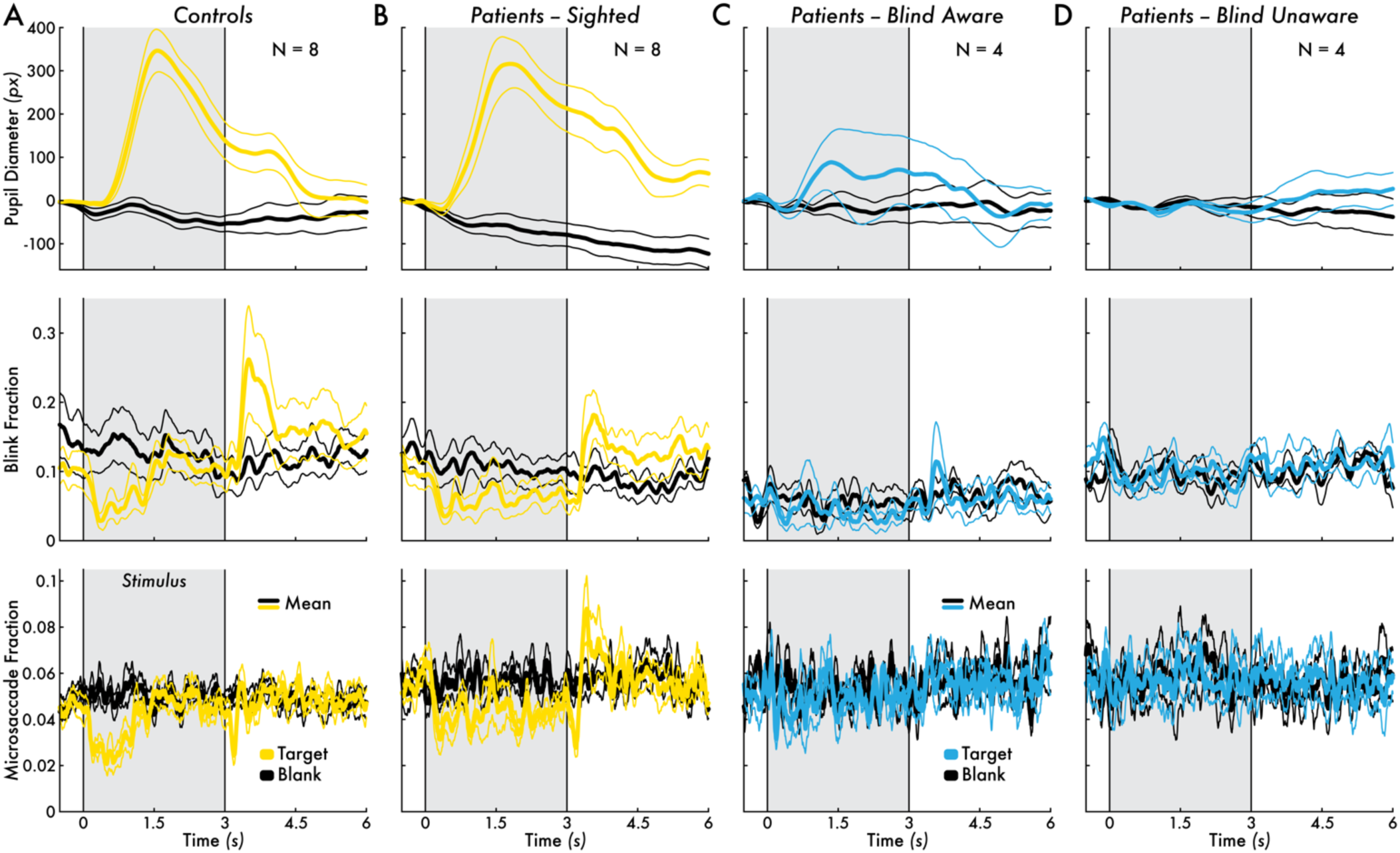
Target stimulus-evoked pupil, blink, and microsaccade dynamics. Pupil diameter, blink fraction, and microsaccade fraction change preceding (0.5 seconds [s]) and following (6 s) the target stimulus for **(A)** control participants (N = 8) averaged between the left and right visual field (yellow), **(B)** patient participants (N = 8) sighted field (yellow), **(C)** blind aware patient participants (N = 4) blind field (blue), and **(D)** blind unaware patient participants (N = 4) blind field. For all subplots, corresponding blank event pupil diameter, blink fraction, and microsaccade fraction responses are shown (black). The group mean eye metric timecourses are shown (thicker traces) bounded by the standard error of the mean (SEM; thinner traces). The 3-second stimulus presentation interval is highlighted by a gray area bounded between two vertical lines at 0 and 3 s.

**Supplementary Figure 4.**
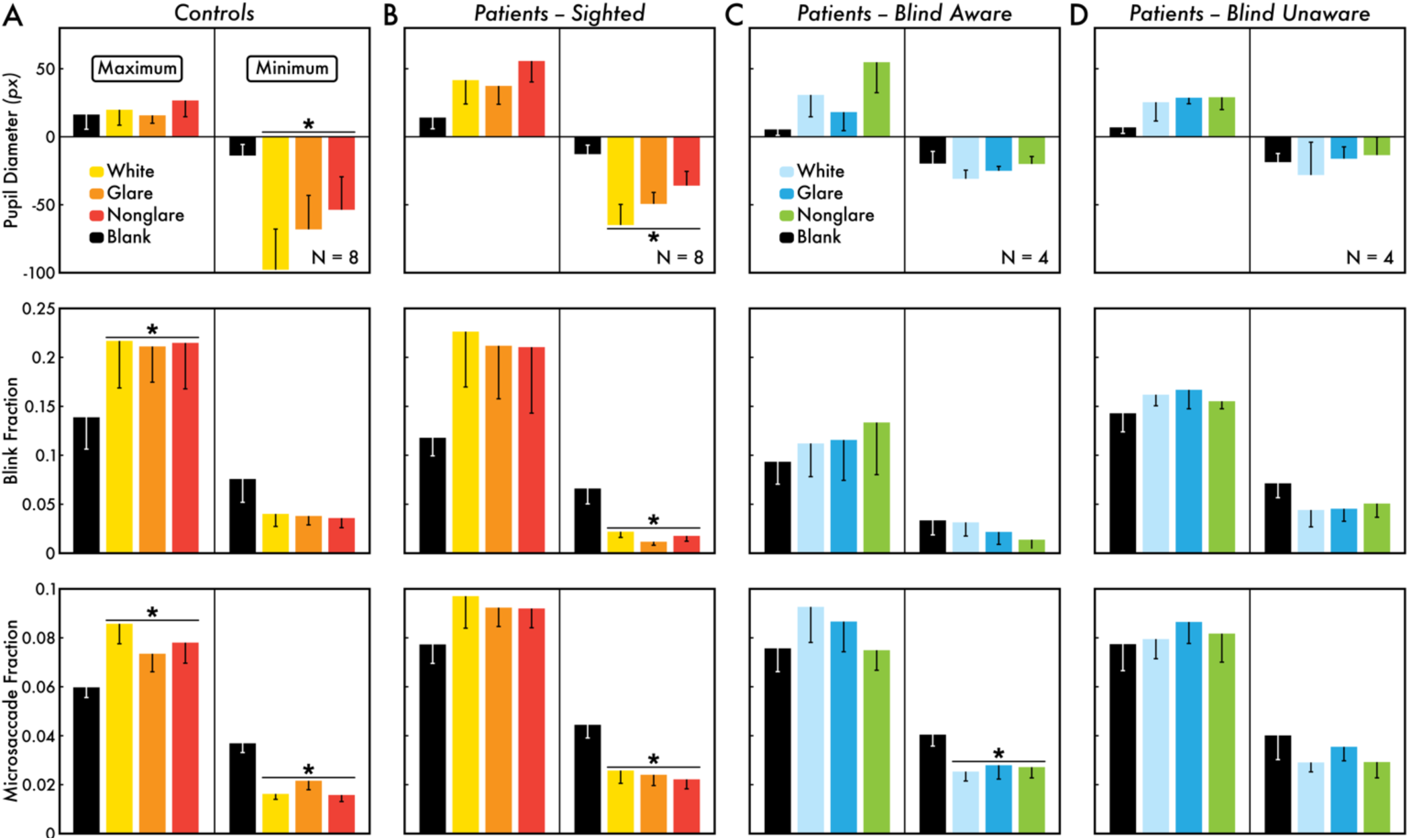
Nontarget stimulus versus blank event pupil, blink, and microsaccade dynamics statistics. Maximum and minimum pupil diameter, blink fraction, and microsaccade fraction change in the first 1.5 seconds (s) following nontarget stimuli (white stimulus = yellow/light blue; glare stimulus = orange/blue; nonglare stimulus = red/green) or blank event (black) for **(A)** control participants (N = 8) averaged between the left and right visual field, **(B)** patient participants (N = 8) sighted field, **(C)** blind aware patient participants (N = 4) blind field, and **(D)** blind unaware patient participants (N = 4) blind field. In all subplots, the bar heights indicate the group averaged maximum and minimum value with error bars showing the standard error of the mean (SEM). Statistically significant (*p* < 0.05) differences of the maximum and minimum change in eye metrics between the nontarget stimuli and blank events are indicated with an asterisk (*).

**Supplementary Figure 5.**
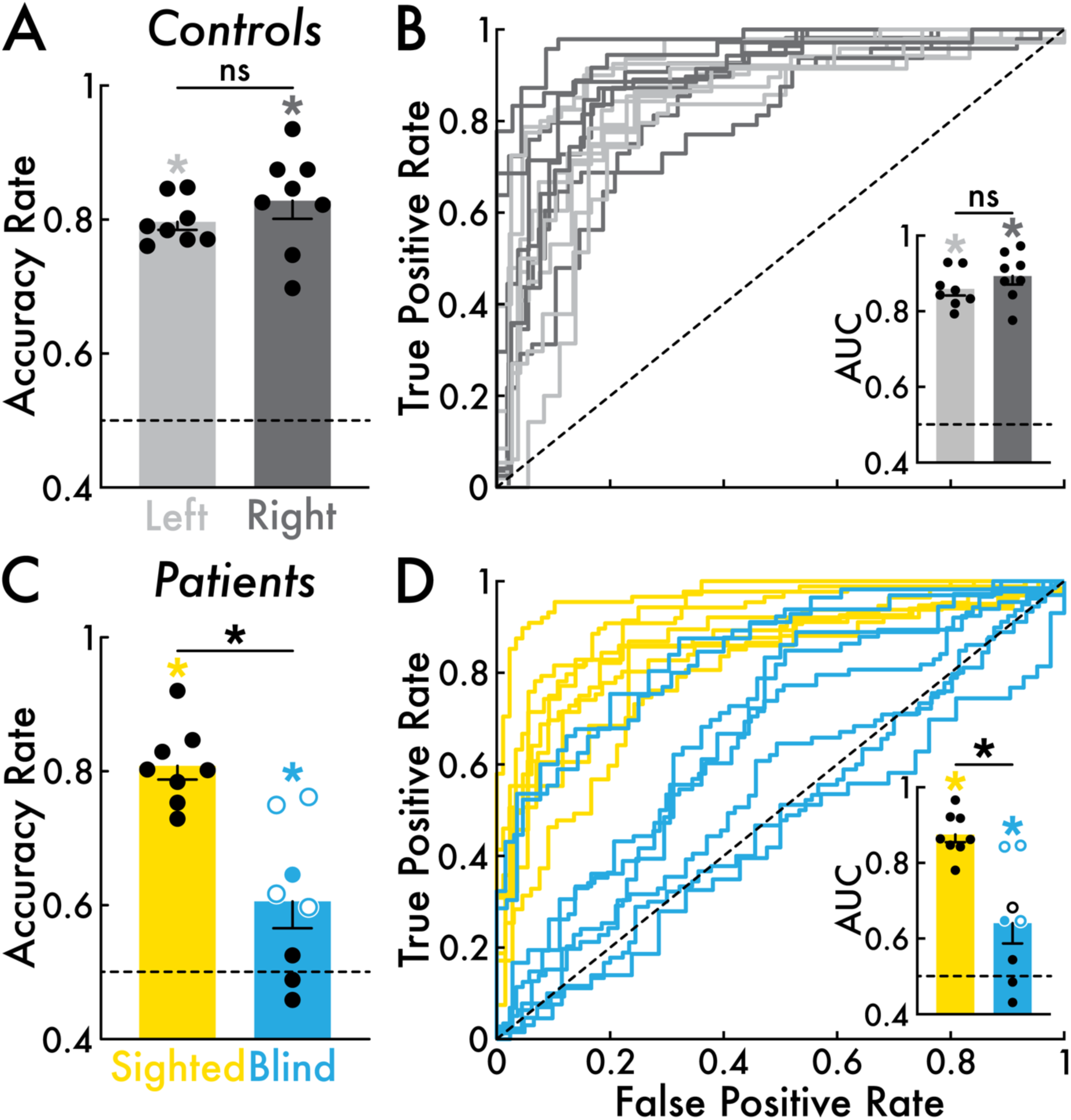
Eye metric-based target stimulus versus blank event classification performance. **(A)** Control participants (N = 8) classification accuracy rate for the left (light gray) and right (dark gray) visual field. Left and right visual field accuracy rates were significantly (*; *p* < 0.05) greater than chance. Left versus right visual field accuracy rates were not significantly different (ns). **(B)** Control participants classification performance visualized by receiver operating characteristic (ROC) curves (false positive rate versus true positive rate). Each timecourse corresponds with a control participant and stimulus presentation location condition (left and right visual field). The inset bar graph depicts the area under the curve (AUC) for each ROC curve. Left and right visual field AUC were significantly (*p* < 0.05) greater than chance. Left versus right visual field AUC was not significantly different. **(C)** Patient participants (N = 8; blind aware patient participants N = 4 [open circles]; blind unaware patient participants N = 4 [closed circles]) classification accuracy rate for the sighted (yellow) and blind field (blue) stimulus presentation locations. Accuracy rate was significantly (*p* < 0.05) greater than chance for the sighted field but not significant for the blind field. Sighted field accuracy rate was significantly (*p* < 0.05) greater than the blind field. Blind field accuracy rate was significantly (*p* < 0.05) greater than chance for four patient participants (highlighted with blue open and closed circles). **(D)** Patient participants classification performance visualized by ROC curves. Each timecourse corresponds with a patient participant and a location condition (sighted and blind field). The inset bar graph depicts the AUC for each ROC curve. AUC (*p* < 0.05) was significantly greater than chance for the sighted and blind field. Sighted field AUC was significantly (*p* < 0.05) greater than the blind field. Blind field AUC was significantly (*p* < 0.05) greater than chance for three patient participants (highlighted with blue open and closed circles). In all subplots, chance level was 0.5 (highlighted with a dotted line), the open and closed circles represent individual participants, and the bars and error bars indicate the group mean and standard error of the mean, respectively.

**Supplementary Figure 6.**
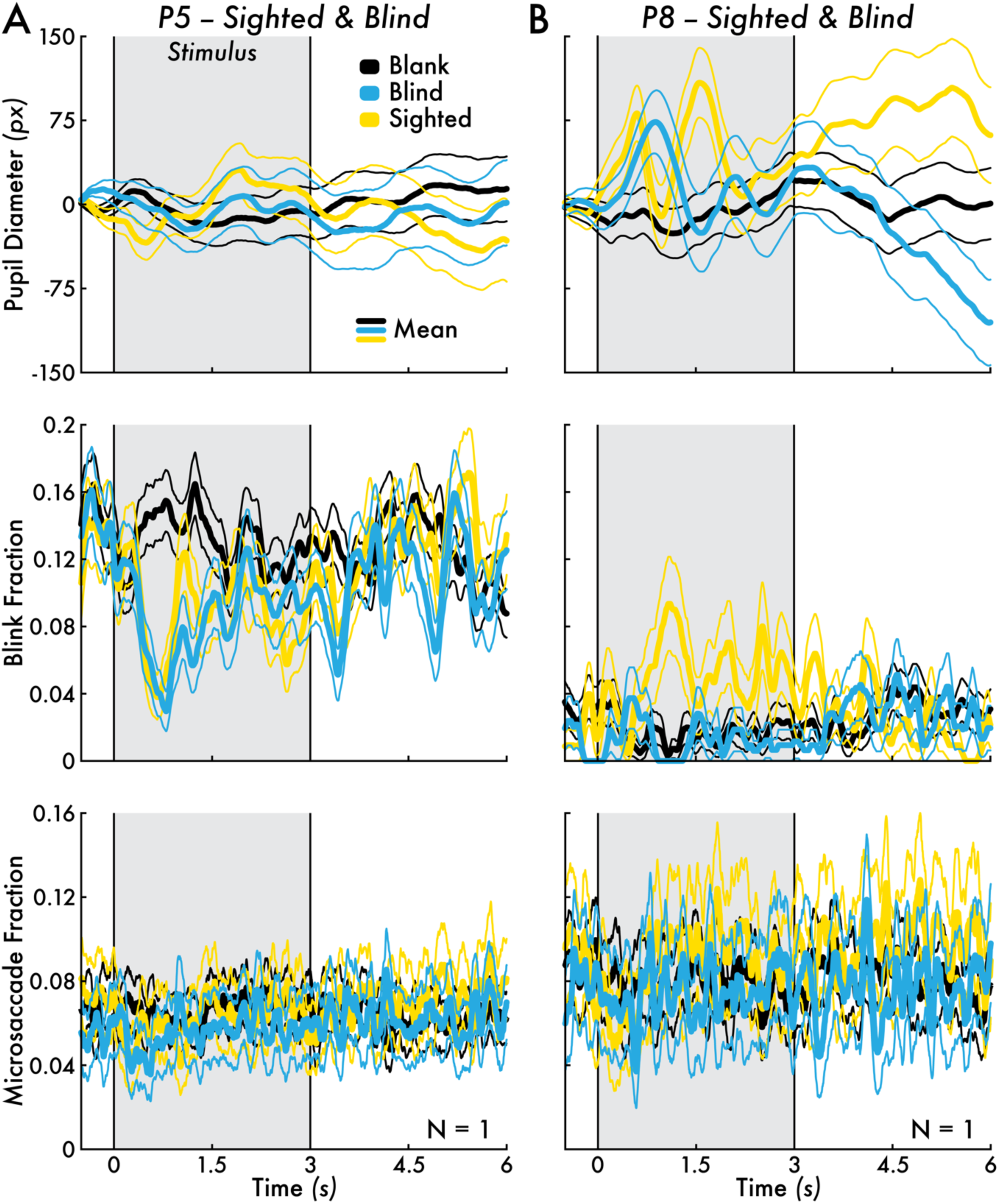
Patient participants P5 and P8 sighted and blind field eye metric dynamics for nontarget stimuli. Patient participant **(A)** P5 and **(B)** P8 sighted (yellow) and blind field (blue) pupil diameter, blink fraction, and microsaccade fraction change for nontarget stimulus versus blank event (black). The mean eye metric changes across all nontarget stimulus trials are shown (thicker traces) bounded by the standard error of the mean (SEM; thinner traces).

**Supplementary Figure 7.**
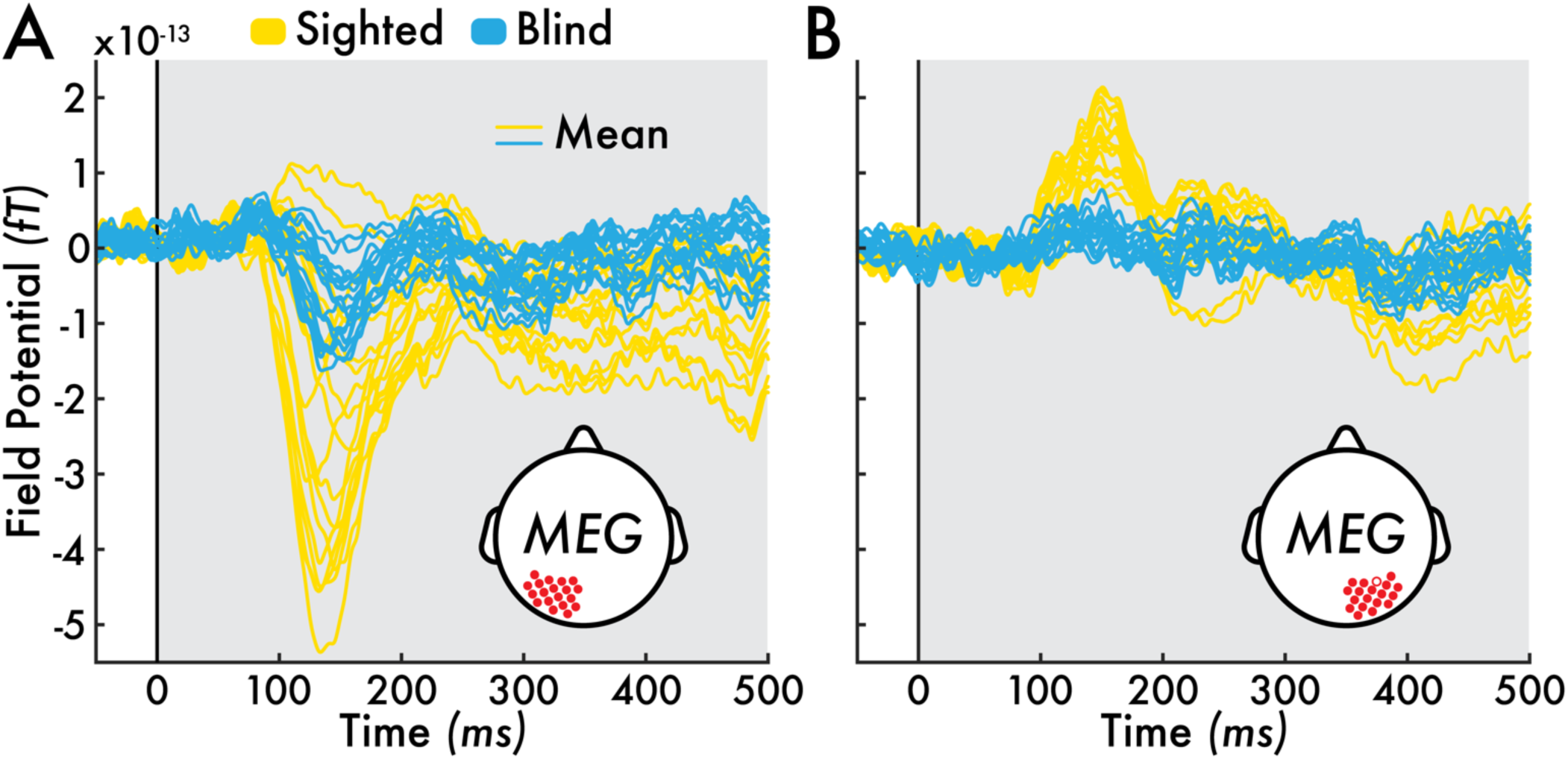
Patient participant P4 sighted and blind field MEG responses for nontarget stimuli. The mean nontarget stimuli evoked magnetencephalography (MEG) field potentials (femtotesla; fT) in the sighted (yellow) and blind field (blue) for **(A)** left (sensors = 19) and **(B)** right occipital sensors (sensors = 18; see inset diagram for approximate sensor locations on the scalp; malfunctioning right occipital sensor O13 was not recorded; highlighted with an open circle). The MEG results were acquired with the MEG-adapted visual perception task (see *Visual Perception Task* Methods section).

